# Reciprocal feature encoding by cortical excitatory and inhibitory neurons

**DOI:** 10.1101/2022.03.14.484357

**Authors:** Adrian J. Duszkiewicz, Pierre Orhan, Sofia Skromne Carrasco, Eleanor H. Brown, Eliott Owczarek, Gilberto R. Vite, Emma R. Wood, Adrien Peyrache

**Affiliations:** Montreal Neurological Institute and Hospital, McGill University; Montreal, QC, Canada; Centre for Discovery Brain Sciences, University of Edinburgh; Edinburgh, UK; Ecole normale supérieure, PSL University, CNRS; Paris, France

## Abstract

In the cortex, the interplay between excitation and inhibition determines the fidelity of neuronal representations. However, while the receptive fields of excitatory neurons are often fine-tuned to the encoded features, the principles governing the tuning of inhibitory neurons are still elusive. We addressed this problem by recording populations of neurons in the postsubiculum (PoSub), a cortical area where the receptive fields of most excitatory neurons correspond to a specific head-direction (HD). In contrast to PoSub-HD cells, the tuning of fast-spiking (FS) cells, the largest class of cortical inhibitory neurons, was broad and heterogeneous. However, we found that PoSub-FS cell tuning curves were often fine-tuned in the spatial frequency domain, which resulted in various radial symmetries in their HD tuning. In addition, recordings and specific optogenetic manipulations of the upstream thalamic populations as well as computational models suggest that this population co-tuning in the frequency domain has a local origin. Together, these findings provide evidence that the resolution of neuronal tuning is an intrinsic property of local cortical networks, shared by both excitatory and inhibitory cell populations.

## Main

Understanding the nature of neural computation is traditionally addressed by determining how external and internal signals are represented at the neuronal level ^1–4^. While neurons in many sensory and other cortical systems encode high-dimensional features ^5–8^, their tuning can only be measured for a limited range of the possible feature space. In comparison, the feature space of the HD system is relatively simple, with excitatory neurons firing for specific directions of the head in the horizontal plane ^4,9,10^. The HD signal is transmitted from the anterodorsal nucleus (ADN) of the thalamus to the cortical recipient neurons in the PoSub ^9–12^. Importantly, this simplicity allows for a full characterization of neuronal tuning during natural behaviors ^13,14^.

Cortical inhibition plays a critical role in shaping the tuning of neuronal responses ^15–23^. Yet, the tuning of inhibitory neurons, especially FS cells, is often considered to be broad ^24–30^ or irregular ^31–35^. By interrogating the HD tuning of PoSub-FS neurons in relation to the tuning of local excitatory cells and upstream thalamic inputs, we show that on the population level PoSub-FS neurons represent features with equal resolution when compared to their local but not upstream excitatory counterparts.

### High-density recordings in the PoSub of freely moving mice

We first established the functional border between PoSub and posterior retrosplenial cortex using a Neuropixel linear electrode array (**Fig. 1a-c**). The step-like increase in average HD tuning along this axis was then used to guide the pre-implanted, microdrive-mounted 64-channel linear electrode arrays and record single-unit activity in PoSub. Probes were implanted either vertically (n = 931 units from 14 mice, range: 46-101 units per recording, **Extended Data Fig. 1a-c**) or parallel to PoSub cell layers (n = 1999 units from 18 mice, range: 42-185 units per recording, **Fig. 1d-e, Extended Data Fig. 1d-e**) and electrode positions were later confirmed histologically (**Extended Data Fig. 1d**). As we found no differences in cell properties across the two datasets (data not shown), they were pooled for further analysis. All recording sessions consisted of square open field exploration and sleep epochs, with a subset of sessions extended to include a triangular open field or a cue rotation task. PoSub units were subdivided into putative excitatory cells (n = 1835) and putative PoSub-FS cells (n = 427) based on mean firing rate and waveform shape (**Extended Data Fig. 2a**). A subset of excitatory cells whose HD information exceeded the 99^th^ percentile of the time-reversed control distribution (see **Methods**) were classified as PoSub-HD cells (> 0.2 bits/spike, n = 1602, 87% of excitatory cells; **Extended Data Fig. 2b**).

**Fig 1.**
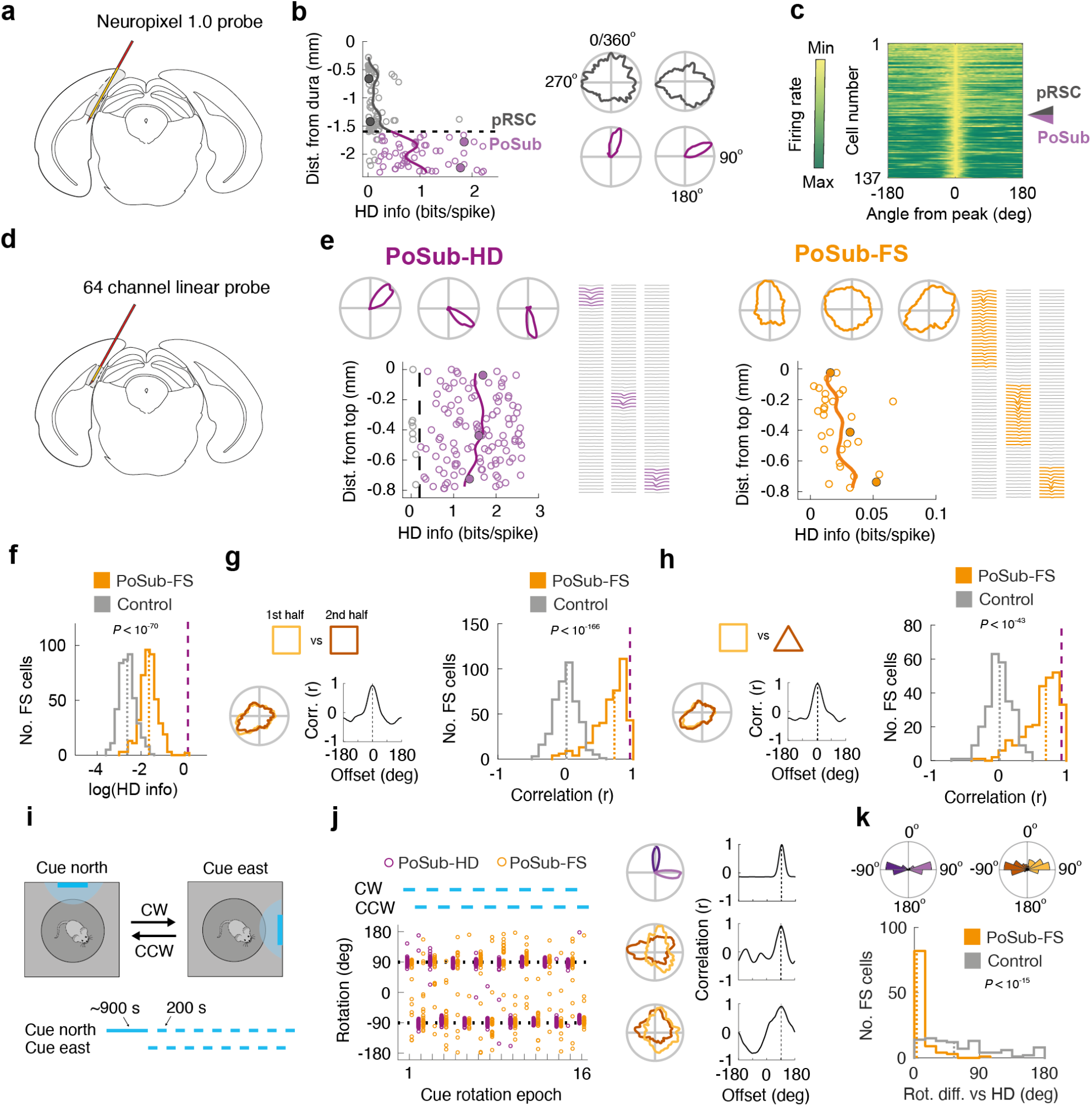
PoSub-FS cells share functional properties with canonical HD cells. **(a)** Brain diagram showing the Neuropixel probe along posterior retrosplenial cortex (pRSC) and PoSub. **(b)** Scatterplot depicting HD information of all putative excitatory cells in a single Neuropixel recording as well as the running average (continuous line). Representative HD tuning curves correspond to filled circles. Dashed line, putative boundary between pRSC and PoSub. **(c)** Normalized tuning curves of all cells in (**b)**, sorted according to the position on the probe. Arrowhead, the putative pRSC/PoSub boundary. **(d)** Brain diagram showing the angled 64-channel linear probe in PoSub. **(e)** Scatterplots depicting HD information of all putative excitatory cells (left) and putative PoSub-FS cells (right) in a single angled recording. Solid lines, running average. Representative HD tuning curves and waveforms correspond to filled circles. **(f)** Histogram of HD information carried by tuning curves of PoSub-FS cells (n = 427; Wilcoxon signed rank test, Z = 17.9, *P* < 10^−70^). Dotted lines, medians of depicted distributions. Dashed lines, median of PoSub-HD cell distribution. (**g-h**) PoSub-FS cell tuning curve correlation across (**g**) two halves of the square arena epoch or **(h)** square and triangle arena epochs. Left: Representative tuning curves of a single PoSub-FS cell and their cross-correlation. Dotted lines, maximum correlation. Right: Population histograms (two halves of square: n = 427, Wilcoxon signed rank test vs time-reversed control, Z = 17.8, *P* < 10^−166^; square vs triangle: n = 264, Wilcoxon signed rank test vs time-reversed control, Z = 14.0, *P* < 10^−44^). Dotted lines show medians of the depicted distributions, dashed lines show the median of the PoSub-HD cell distribution. **(i)** Cue rotation task. Top: diagram of the cue rotation apparatus. The mouse was placed on a small elevated platform in a dim recording chamber, with the light from the distal cue providing the only light source. The cue was then rotated back-and-forth either clockwise (CW) or counterclockwise (CCW) by 90° every 200 seconds. Bottom: timeline of the epochs corresponding to different cue positions. **(j)** Representative cue rotation session. Left: Rotation of PoSub-HD and PoSub-FS tuning curves over 16 consecutive cue rotation epochs (blue lines). Each point denotes tuning curve rotation of a single cell relative to the previous epoch. Right: representative PoSub-HD and PoSub-FS tuning curves from the same session computed across all CW and CCW epochs (light and dark shades, respectively), and their cross-correlation. Dotted line, maximum correlation. **(k)** Population data from the cue rotation task. Top: Distribution of average tuning rotations of PoSub-HD cells (n = 411) and PoSub-FS cells (n = 99) across all CW and CCW epochs (light and dark shades, respectively). Bottom: Histogram of mean absolute rotation differences between individual PoSub-FS cells and the average of PoSub-HD cells (n = 99, Wilcoxon signed rank test vs time-reversed control, Z = 8.25, *P* < 10^−15^). Dotted lines, medians of the depicted distributions.

### PoSub-FS cells share functional properties with canonical HD cells

At the population level, we observed that the quantity of HD information conveyed by PoSub-FS cells was significantly higher than that of the time-reversed control population (**Fig. 1f**). However, in contrast to canonical HD cells, individual PoSub-FS cells had complex, often multi-peaked tuning curves not confined to discrete HD values. We hypothesized that since PoSub-FS cells receive inputs from local PoSub-HD cells (**Extended Data Fig. 2c-d**), they would share their functional properties. Indeed, the tuning of PoSub-FS cells was stable within a single exploration epoch (**Fig. 1g)** and independent of the enclosure geometry (**Fig. 1h**), reflecting the properties of canonical PoSub-HD cells (**Fig. 2e-f**). Importantly, in a cue rotation paradigm ^36,37^ (**Fig. 1i, Extended Data Fig. 2g**) PoSub-FS cell tuning followed the rotated distal landmark in concert with PoSub-HD cells (**Fig. 1j-k**). Thus, while PoSub-FS cells, in contrast to canonical HD cells, exhibit irregular HD tuning curves, they are functionally integrated into the cortical HD system.

**Fig 2.**
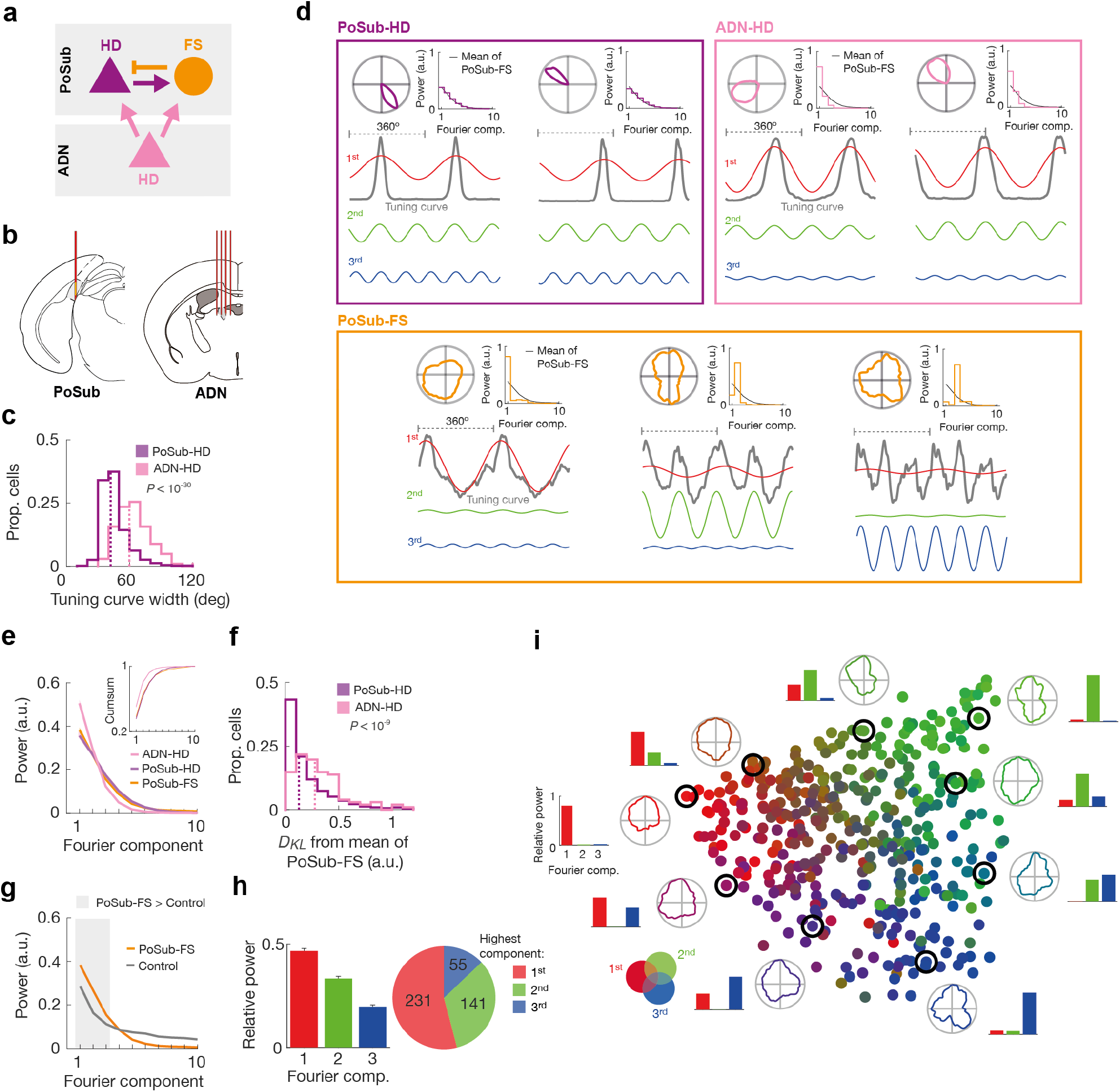
HD tuning of PoSub-HD and PoSub-FS cell populations is equivalent in the Fourier space. **(a)** Simplified diagram of connections in the thalamocortical HD system. **(b)** Brain diagram showing probes in PoSub and the anterior thalamus. **(c)** Histograms showing tuning curve widths of PoSub-HD and ADN-HD cells (n = (1602, 97), Mann-Whitney U test, Z = 11.4, *P* < 10^−30^). Dotted lines, medians of the depicted distributions. **(d)** Fourier decomposition of representative PoSub-HD, ADN-HD and PoSub-FS tuning curves. Each example depicts the tuning curve in polar coordinates (top left), normalized power of first ten Fourier components (top right), and two periods of linearized tuning curve along with first three Fourier components in sine wave form. Black curves in Fourier signatures, average PoSub-FS spectrum. **(e)** Average Fourier signatures of PoSub-FS, PoSub-HD and ADN-HD cell populations (2-way ANOVA, Fourier component by cell type interaction: F_(9,19116)_ = 2.76, *P* < 0.01). Inset: cumulative distribution of the data in the main panel. Shaded area, SEM. **(f)** Statistical distance between individual PoSub-HD cell or ADN-HD cell Fourier spectra and the average Fourier signature of the PoSub-FS cell population (n = (1602, 97), Mann-Whitney U test, *P* < 10^−9^). Dotted lines, medians of the depicted distributions. *D*_*KL*_, Kullback-Leibler divergence. **(g)** Average Fourier signature of PoSub-FS cells compared with time-reversed control (n = 427, 2-way ANOVA, Fourier component by cell type interaction: F_(9,3834)_ = 64.1, *P* < 10^−37^). Gray background represents pairwise comparisons in which PoSub-FS group is significantly higher than control group (Wilcoxon signed rank test with Bonferroni correction, all *P* < 10^−4^). Shaded area, SEM. **(h)** Right, relative mean power of the first three Fourier components for PoSub-FS cells. Right, proportions of PoSub-FS cells with the highest power in the each of the first three Fourier components. Error bars, SEM. **(i)** Two-dimensional Isomap projection of PoSub-FS cell tuning curve auto-correlograms. Points represent auto-correlograms of individual PoSub-FS cells, coloured using the red-green-blue (RGB) colour model mapped to the relative power of the first three Fourier components. The projection is surrounded by representative PoSub-FS cell tuning curves and the relative power of the first three Fourier components. Black circles, points corresponding to the adjacent representative cells.

### HD tuning of PoSub-HD and PoSub-FS cells is equivalent in the Fourier space

We next aimed to establish whether the tuning of PoSub-FS cells was related to the tuning of local HD cells. To address this question, we compared the HD tuning of PoSub-FS cells with that of PoSub-HD cells as well as with that of the HD cells in the upstream ADN (ADN-HD cells; n = 97 cells from 8 mice; **Fig. 2a-b**). In mice, ADN-HD neurons tend to have broader HD tuning curves than PoSub-HD neurons (**Fig. 2c, Extended Data Fig. 3a-b**), a property independent of their tendency to fire in anticipation of future HD ^38^ (**Extended Data Fig. 3c-d**). We thus utilized these differences in tuning curve shape between the two populations to characterize their relative influence on the tuning of PoSub-FS cells. Importantly, HD tuning of PoSub-FS cells cannot be directly compared with that of canonical HD cells due to their irregular, often multi-peaked shape. In order to overcome this, we transformed the HD tuning curves from the spatial domain (“HD space”) to the spatial frequency domain (“Fourier space”). (**Fig. 2d**; see **Methods**). Each tuning curve was thus represented as a sum of sine waves (“Fourier components”) whose frequencies are equal to the harmonics of the unit circle, corresponding to periods from 360° (fundamental frequency) to 2° (highest possible harmonic, equal to twice the tuning curve sampling bin). In turn, each Fourier component could be described in terms of its amplitude (or “power”) and phase, which reflects the relative orientation of that component. Each tuning curve was thus associated with an individual “Fourier signature”, consisting of relative powers of its Fourier components. Across the cell population, the Fourier power decayed rapidly as a function of frequency. Hence, for clarity, we focused our analysis on the relative power of the first ten Fourier components which contained, on average, 98% of the total power.

Canonical HD cells, sharply tuned in HD space, showed broad and stereotyped tuning in Fourier space, with power distributed across several Fourier components and each successive component showing progressively less power. We found that Fourier signatures of PoSub-HD and ADN-HD cells often differed between the two regions (**Fig. 2d**, top): ADN-HD Fourier signatures tended to be skewed towards lower components compared to PoSub-HD cells, reflecting their broader tuning curves.

In contrast, PoSub-FS cells exhibited highly heterogenous Fourier signatures, with many cells broadly tuned in HD space but narrowly tuned in the Fourier space, i.e. showing high power for only one Fourier component (**Fig. 2d**, bottom). We hypothesized that while Fourier signatures of individual PoSub-FS cells reflect their heterogenous tuning shapes, overall they are ought to be constrained by the tuning properties of their main HD inputs. Indeed, we found that the average Fourier signature of PoSub-FS cells was similar to the Fourier signature of PoSub-HD cells but unlike that of ADN-HD cells (**Fig. 2e-f, Extended Data Fig. 3f**). Importantly, the mean Fourier signature of PoSub-FS cells was stable across different environmental manipulations (**Extended Data Fig. 3n**), reflecting the stability of their tuning in spatial domain. Like hippocampal place cells, HD cells can sometimes have multiple receptive fields ^34^ which could affect the average Fourier signature of an HD cell population. However, the reported similarity between PoSub-HD and PoSub-FS cell Fourier signatures was even more pronounced when the dataset was limited to HD cells with a single receptive field (**Extended Data Fig. 3h-l**). Thus, although HD tuning curves of individual PoSub-HD and PoSub-FS cells appear strikingly different, on a population level they share the same underlying Fourier power spectrum.

Interestingly, all Fourier modes were not equally important to explain the tuning of the PoSub population. Compared to controls, the spectrum of PoSub-FS cells showed higher power for the first three Fourier components only (**Fig. 2g**), suggesting that these three axes of symmetry form the tenet of directional tuning in the PoSub. We thus used the first three Fourier components to classify cells according to their most prominent axis of symmetry. The proportions of recorded PoSub-FS cells with the highest relative power in each of the first three Fourier components were similar to the relative mean power for the population (**Fig 2h**).

Although they shared the same average spectrum, HD and FS cell populations in the PoSub differed in two main aspects. First, while the shape of the Fourier signature was largely uniform across HD cells from the same brain region, it was highly variable across PoSub-FS cells (**Extended Data Fig. 3g**), indicating narrow tuning of PoSub-FS cells in the Fourier space (**Fig. 2d**). Hence, the Fourier spectrum of the local HD signal was distributed among the PoSub-FS cell population rather than being homogeneously reflected within each individual cell, resulting in symmetrical tuning curves. Second, in individual HD cells, the phases of Fourier components were correlated with each other, as expected for any symmetrical function with a single maximum (**Extended Data Fig. 3m**, *left* and *middle*). In contrast, the Fourier components of individual PoSub-FS cells had random phases relative to each other (**Extended Data Fig. 3m**, *right*), explaining the apparent irregularity of their HD tuning for PoSub-FS cells showing power in more than one Fourier component.

To gain further insights into the shape of PoSub-FS cell tuning curves independently of their relative orientation, we computed an auto-correlation function for each cell by correlating its tuning curve with itself at different circular offsets. We then projected the resulting PoSub-FS cell auto-correlograms onto a 2-dimensional space using the Isomap dimensionality reduction algorithm ^39^ (**Fig. 2i, Extended Data Fig. 4a**). The resulting projection reflects the heterogeneity of PoSub-FS cell tuning curve shapes across the population (compared to control data, **Extended Data Fig. 4b**). Importantly, the triangular shape of this unsupervised embedding revealed that a large portion of the power was concentrated in the first three Fourier components. PoSub-FS cells located at each vertex of the triangle showed pure 1-, 2-, or 3-fold symmetrical HD tuning (**Fig. 2i, Extended Data Fig. 4a)**, reflecting their narrow tuning in the Fourier space. Still, this distribution was a continuum and other PoSub-FS cells were characterized by a subset of Fourier components (**Fig. 2i**). We then projected the data again into the same 2-dimensional space, this time adding the auto-correlograms of PoSub-HD and ADN-HD cells. We found that PoSub-HD and ADN-HD cell auto-correlograms occupy compact subspaces within the broader distribution, reflecting their relative homogeneity (**Extended Data Fig. 4c**). The auto-correlogram distribution of PoSub-HD cells occupied the centre of PoSub-FS cell distribution while that of ADN-HD cells was closer to the periphery (**Extended Data Fig. 4d**), confirming the observations of differences in average Fourier signatures (**Fig. 2e-f**). In conclusion, the shapes of PoSub-FS cell tuning curves were broadly distributed and each was unique. Yet, their tuning resolution was shared with PoSub-HD cells, but not ADN-HD cells.

### PoSub-FS cell tuning reveals key circuit properties

To account for the origin of PoSub-FS cell tuning, we turned to numeric simulations and theory. First, we computed the tuning of output units in a fully connected network receiving HD-tuned inputs (**Fig. 3a-c**). Output neurons linearly integrated their inputs, as it is the case for cortical FS cells ^40^. The spectrum of input tuning curves directly depended on the tuning curve width (**Fig. 3a**). Output neurons can show symmetries similar to those observed in PoSub-FS neurons (**Fig. 3b**). Importantly, although the input and output tuning curves were strikingly different, random connectivity preserved the mean Fourier spectrum of tuning (**Fig. 3c**).

**Fig 3.**
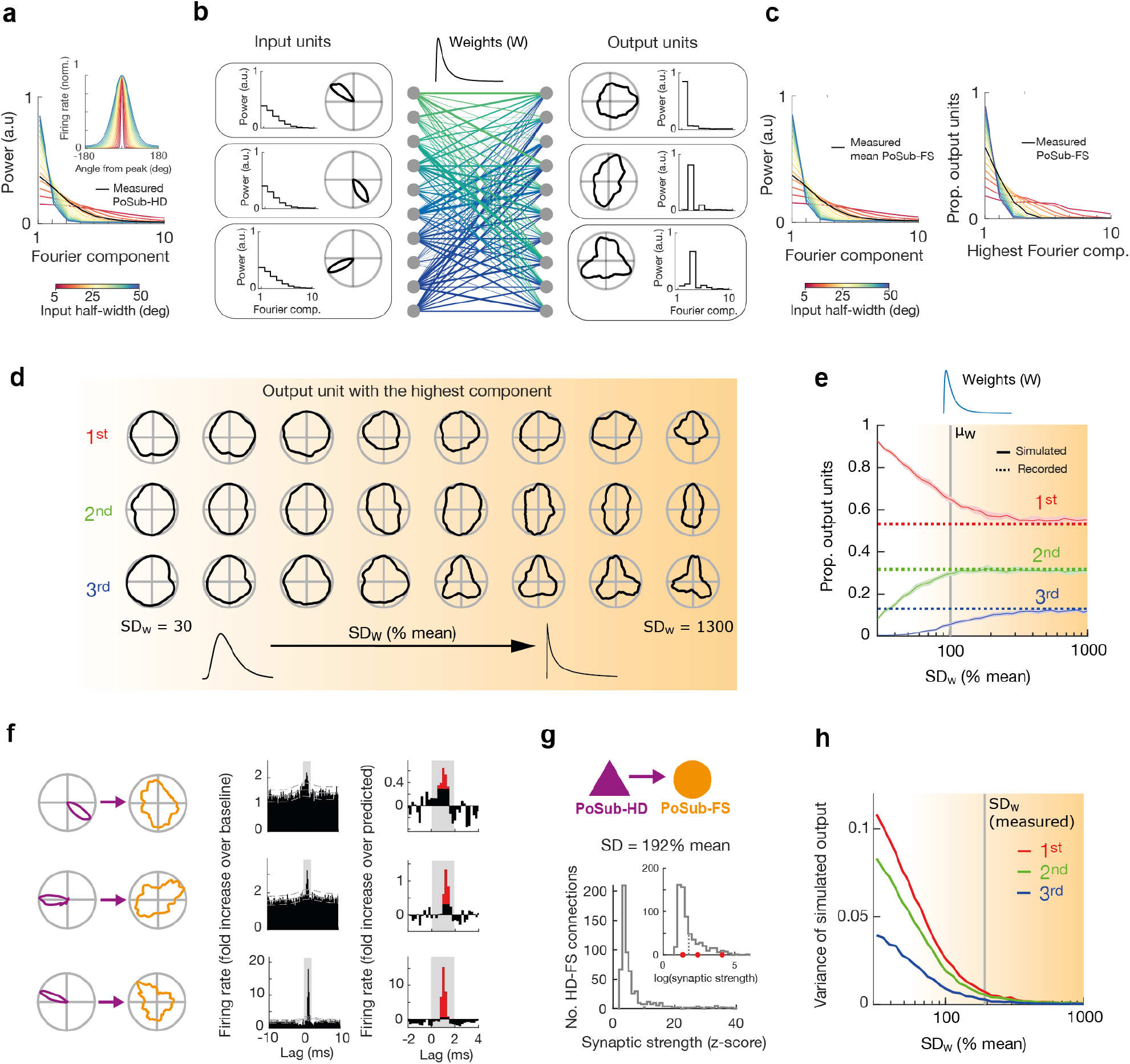
PoSub-FS like tuning curves emerge from random connectivity in a linear regime and reveal circuit properties. (**a**-**c**) Input and output units share the same average Fourier signature in simulation. **(a)** Relationship between tuning curve width and Fourier signature of simulated HD-like input tuning curves. **(b)** Emergence of symmetries from random feed-forward connections. Left, example HD cell tuning curves recorded in PoSub. Middle, random connectivity with log-normal distribution. Right, tuning curves resulting from random linear combinations of input HD tuning curves. **(c)** Left, the mean Fourier power of the normalized random linear combinations of canonical input tuning curves. Right, ratios of output tuning curves with highest power in each Fourier component as a function of input tuning curve width. **(d)** Output tuning curves with highest Fourier power in the first three components as a function of spread of weight distribution in an example simulation. The width of weight distribution is expressed as standard deviation (SD) divided by mean. **(e)** Proportion of simulated cells with maximum power in the first three Fourier component as a function of spread of synaptic weight distribution. Dotted lines display actual proportion of recorded PoSub-FS cells. Shaded area of each curve, SD based on 40 simulations. (**f-g**) Estimating variance in synaptic strength from putative excitatory connections. **(f)** Example tuning curves of putative synaptic HD-FS cell pairs (left), spike cross-correlations at high resolution (0.2 ms bins, middle) and high magnifications of putative synaptic peaks (right). Red bars, portion of the cross-correlogram significantly higher than predicted. **(g)** Histogram of Z-scored putative synaptic strength (amplitude of the short-latency peak). Inset: logarithm of the same variable. Red dots, values corresponding to the examples in F. Dotted lines, medians of the depicted distributions **(h)** Variance in proportion of simulated cells with maximum power at each component (over 30 simulations). Simulations consistently result in proportions close to the observed values when the spread in synaptic weight distribution exceeds the value determined experimentally.

To reveal the conditions under which random connectivity leads to symmetrical tuning similar to the real data, we varied two features of the network: the variance of weight distribution and the number of input units. In simulation, we observed that for low variance of the weight distribution, and quite intuitively, all output neurons in the network showed similar tuning curves (**Fig 3d**). When the variance of the input weights was increased, individual output tuning curves progressively showed differences in tuning. This was independent of the specific shape of the distribution of weights (**Extended Data Fig. 5a**). As a result, the proportion of output units showing maximal power in one of the first three Fourier components progressively converged to the proportions observed in PoSub-FS cells (**Fig. 3e, Extended Data Fig. 5b-d**). This convergence emerged for weight distributions in which the standard deviation exceeded the mean of the weights. To test whether this predicted how PoSub-HD cells were connected to PoSub-FS cells, we quantified the strength of putative synaptic connections from spike train cross-correlation of PoSub-HD:PoSub-FS cell pairs ^41^ (**Fig. 3f**) (see **Methods**). The distribution was heavy-tailed (**Fig. 3g**), with standard deviation exceeding the average weight (ratio = 192 %), in the range where *in silico* simulation predicted symmetries to emerge (**Fig. 3h**).

Varying the parameters of the weight distribution did not account for the observed amount of HD information conveyed by PoSub-FS cells (**Fig. 1f**). Rather, we found that the number of input units received by each output unit was a key factor influencing the amount of HD information (**Extended Data Fig. 5e**). Varying both weight distribution and the number of input units, we obtained a distribution of HD information in output tuning curves that matched the real data (**Extended Data Fig. 5f**), revealing that the tuning of PoSub-FS cells can be used to estimate both the distribution of weights and the number of input neurons. Importantly, under optimal network conditions, Isomap projection of output tuning curve autocorrelograms has a similar geometry to that of real PoSub-FS cells (**Extended Data Fig. 5g**), confirming similar distribution of tuning shapes.

Finally, we demonstrated that for linearly integrating units, the output tuning curves of a randomly connected network with high weight variance each have a unique spectrum in which Fourier components are independent and normally distributed (see Methods). Importantly, the mean of the output spectrum is entirely defined by the input tuning curve spectrum while the variance depends on the weight variance. As a result, for high synaptic weight variance, many output neurons show high power in only one Fourier component (and thus striking radial symmetries in the tuning curves), while one the population level, the spectra of input and output units are still similar, as observed in the data.

### PoSub-FS cells receive directionally uniform input from the anterior thalamus

Thalamocortical neurons exert a strong excitatory drive onto FS cells in many cortical areas ^42,43^, including in the ADN-PoSub circuit ^44^. To determine whether upstream thalamic inputs shape PoSub-FS cell tuning, we selectively manipulated the strength (or “gain”) of the thalamic input from ADN to PoSub and quantified the effect of this manipulation on the tuning of PoSub-FS cells. We reasoned that if each PoSub-FS cell receives non-uniform thalamic HD input, increasing input gain should result in non-uniform (multiplicative) modulation of their HD tuning. In contrast, if the thalamic input is uniform, PoSub-FS cell tuning should be uniformly (additively) modulated ^45^.

The ADN is strongly innervated by inhibitory afferents from the thalamic reticular nucleus (TRN, **Extended Data Fig. 6**) ^46,47^. We leveraged this specific inhibitory pathway to selectively increase the activity of ADN-HD cells. To that end, we injected a Cre-dependent AAV-ArchT into TRN of VGAT-Cre mice and recorded ensembles of anterior thalamic neurons (**Extended Data Figure 7a**; n = 127 thalamic cells, including 52 HD cells, from 3 mice). Targeted illumination of ADN (thus inactivating the inhibitory presynaptic terminals of the TRN neurons) resulted in a net increase in the firing rates of ADN-HD cells but not that of not directionally-tuned neurons (non-HD cells) recorded during the same sessions (**Extended Data Fig. 7b-c**), further confirming preferential TRN innervation of ADN over other anterior thalamic nuclei (**Extended Data Fig. 6a-b**). Contrary to long-term disinhibition ^47^, short-term disinhibition of ADN-HD cells was not associated with broadening of their tuning curves (**Extended Data Fig. 6d**), This manipulation therefore selectively increased the gain of the thalamic HD signal without affecting its resolution. We thus used this method to characterize the effects of thalamic gain modulation on the HD tuning of PoSub neurons. We recorded ensembles of PoSub neurons in VGAT-Cre mice injected with AAV-ArchT (n = 83 PoSub-HD cells, 47 PoSub-FS cells from 5 mice) or a control viral vector (n = 89 PoSub-HD cells, 38 PoSub-FS cells from 3 mice; **Fig. 4a**) bilaterally into TRN. Similarly to the upstream ADN-HD cells, PoSub-HD and PoSub-FS cells increased their firing rates (**Fig. 4b, Extended Data Fig. 7e-f**) without significantly distorting their HD tuning (**Extended Data Fig. 7g-j**).

**Fig 4:**
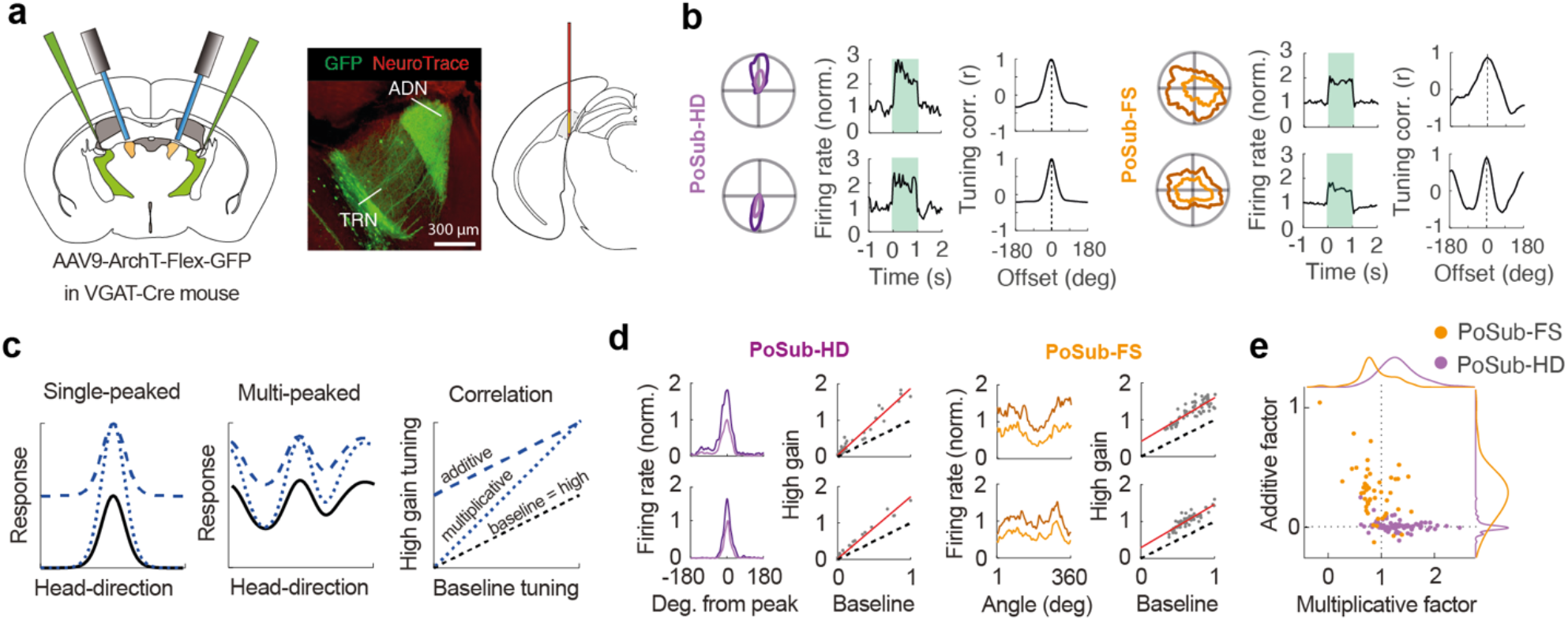
Thalamic drive provides uniform HD input to PoSub-FS cells. **(a)** Experimental setup. Left: Brain diagram of bilateral AAV injection into TRN and bilateral optic fibers above ADN. Middle: Image of a coronal section from the anterior thalamus extensive inhibitory projections from TRN to ADN. Right: Brain diagram of probe placement in PoSub. **(b)** Representative examples of (top) PoSub-HD cell responses and (bottom) PoSub-FS cell responses to the optogenetic elevation of thalamic HD gain. Left: HD tuning curves for the light OFF epoch (light shade) and the light ON epoch (dark shade). Middle: Effect of the optogenetic manipulation on representative cells’ firing rates. Green shading, duration of the light pulse. Right: cross-correlation between tuning curves during the light ON and light OFF epochs. **(c)** Illustration of additive and multiplicative effects of gain modulation on (left) single-peaked and (middle) multi-peaked tuning curves. Right: correlation between HD tuning curves in baseline conditions (black lines) and high gain conditions (blue lines) reveals the contribution of additive and/or multiplicative factors. **(d)** Representative examples of PoSub-HD cells (left) and PoSub-FS cells (right) in baseline condition (light shades) and high gain condition (dark shades) plotted in Cartesian coordinates (same examples as in **b**) as well as their respective tuning correlation plots. Red lines represent the linear fit. **(e)** Scatter plot showing the values of additive and multiplicative factors for each PoSub-HD and PoSub-FS cell as well as normalized population distributions.

We then computed, for each cell, linear regression between HD tuning curves in the baseline condition and under high thalamic gain (i.e. ADN disinhibition). The slope of the linear fit denotes the multiplicative modulation of tuning by thalamic gain, while the intercept denotes additive modulation ^45^ (**Fig. 4c**). Thus, a slope above 1 indicates the presence of multiplicative gain and a positive intercept indicates the presence of additive gain. When computing the tuning curves, we used larger angular bins (6°) instead of Gaussian smoothing in order to ensure that individual points on the tuning curve are independent from one another. We then assessed the contribution of these additive and multiplicative factors to the tuning modulation of PoSub-HD and PoSub-FS cells. The modulation of PoSub-HD cells was purely multiplicative (**Fig. 4d-e, Extended Data Fig. 8A**), indicating that they receive HD-specific thalamic inputs. Indeed, this high degree of multiplicative modulation largely reflected the modulation of the upstream ADN-HD cells (**Extended Data Fig. 8c-e**). In contrast, the modulation of PoSub-FS cell tuning was exclusively additive (**Fig. 4d-e, Extended Data Fig. 8b**), indicating that the thalamic inputs received by individual PoSub-FS cells are uniform across across all directions (**Extended Data Fig. 8f**).

### PoSub-FS cell tuning arises from coupling to the HD ring manifold

Finally, in order to exclude the possibility that the tuning of PoSub-FS cells is determined by external factors like vision, we sought to establish whether their activity is coupled to the internal attractor dynamics in absence of sensory input. The HD signal in the ADN-PoSub pathway is coherently organized into a 1-dimentional (1-D) ring attractor even during sleep when sensory inputs are virtually absent ^11,48^. We thus tested whether the tuning of PoSub-FS cells relies on the intrinsic dynamics of the HD cell attractor network during sleep. To address this question, we analyzed ensemble activity during Rapid Eye Movement (REM) sleep, when the coordination of PoSub-HD cells is virtually indistinguishable from wakefulness ^11^ (**Extended Data Fig. 9a-b**).

We first sought to establish whether the temporal coupling between individual PoSub-FS and PoSub-HD cells was preserved during REM sleep. In order to account for the coupling to the population firing rate irrespective of any specific tuning, we quantified the pairwise coupling between PoSub cells using a General Linear Model (GLM) ^49^. While both PoSub-HD and PoSub-FS cells showed strong coupling to the population (**Extended Data Fig. 9c**), we found that the polarity of the GLM cross-coupling coefficient between PoSub-HD cell pairs was preserved across wakefulness (WAKE) and Rapid Eye Movement (REM) sleep (**Fig. 5a-c**, top). For example, PoSub-HD cell pairs co-active during wakefulness showed a high degree of co-activity during REM sleep, while those that were negatively coupled during WAKE were also negatively coupled during REM. Similarly, PoSub-FS cell pairs also preserved their coupling across WAKE and REM, albeit to a smaller degree than PoSub-HD cell pairs (**Fig. 5a-c**, middle). Importantly, the two cell populations were also coupled to each other across WAKE and REM (**Fig. 5a-c**, bottom). Predictably, for all cell pairs the polarity of the cross-coupling coefficient depended on the HD tuning relationship within each cell pair during both WAKE and REM (**Extended Data Fig. 9d**). Overall, these results indicate that the activity of PoSub-FS cells is coupled to the internal attractor dynamics of the HD system.

**Fig 5.**
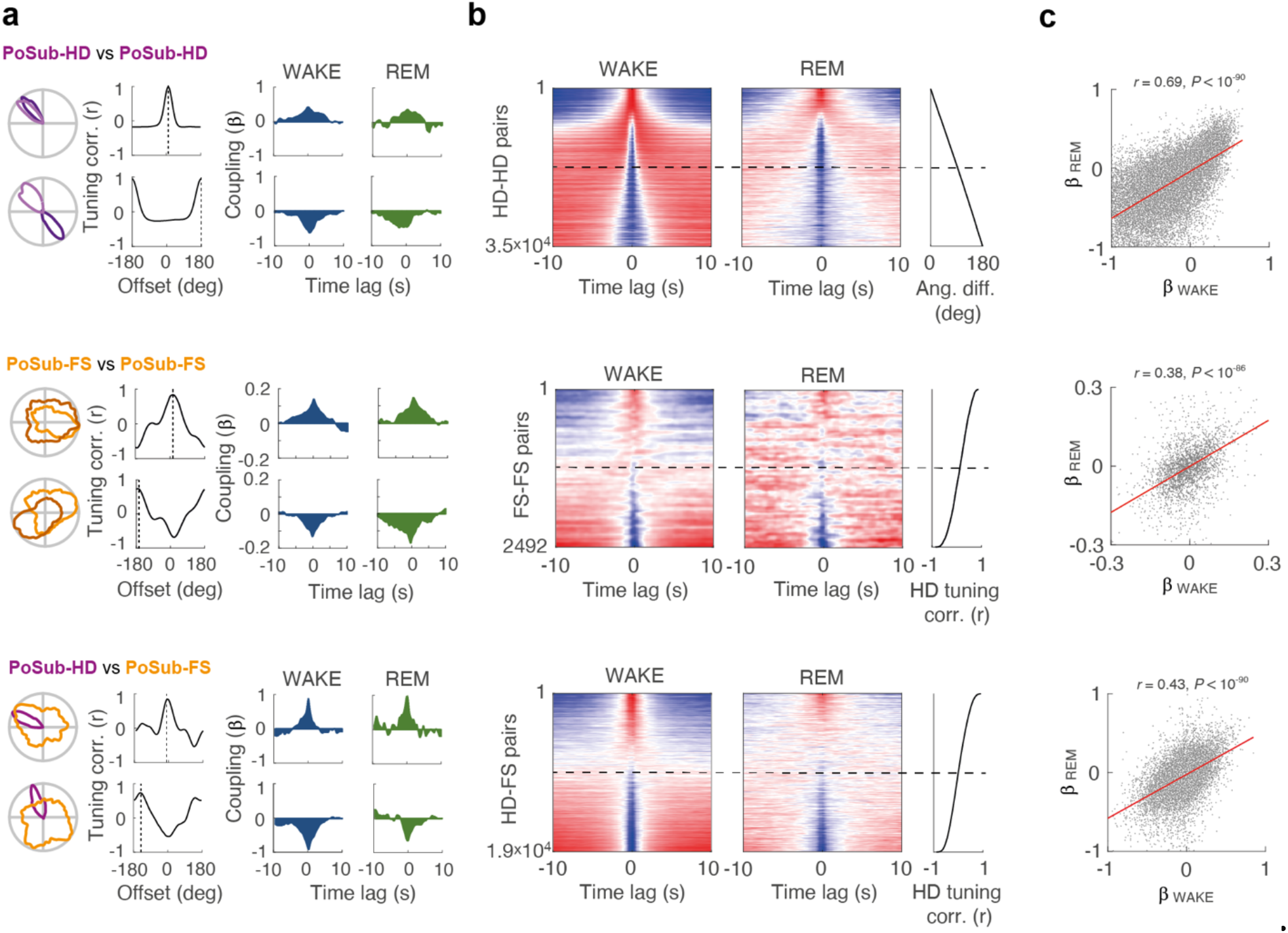
PoSub-HD and PoSub-FS cells show conserved dynamics across WAKE and REM. (**a-c**) Cross-coupling relationships between PoSub-HD:PoSub-HD cell pairs (top row), PoSub-HD:PoSub-FS cell pairs (middle row) and PoSub-FS:PoSub-FS cell pairs (bottom row). **(d)** HD tuning and spike-timing relationships of representative cell pairs. Left: Superimposed HD tuning curves and their HD tuning cross-correlation. Dotted line shows the offset of maximum correlation. Right: GLM cross-coupling during WAKE and REM. **(e)** Color-mapped GLM cross-coupling of all cell pairs during WAKE (left) and REM (middle). Cell pairs were sorted (right) according to the angular difference of their tuning curves (for PoSub-HD:PoSub-HD pairs) or tuning curve correlation at zero offset (PoSub-HD:PoSub-FS and PoSub-FS:PoSub-FS pairs). Each row represents a normalized cross-coupling curve of a single cell pair, color-mapped from minimum (blue) to maximum (red). **(f)** Scatter plots illustrating the cross-coupling correlation across WAKE and REM. Red line represents the linear fit. β, cross-coupling coefficient.

While stable correlation structures among cell pairs constitute strong evidence for coupling of PoSub-FS cells to the HD attractor network, large-scale population recordings enable a more direct visualization and analysis of the 1-D ring attractor manifold that constrains the activity of individual HD cells. We thus asked whether the activity of PoSub-FS cells is constrained by the same ring attractor manifold as that of PoSub-HD cells. To that end, we applied Isomap ^39^ to HD cell population vectors in order to visualize the 1-D ring manifold of PoSub-HD cell population activity during WAKE ^48,50^ (**Fig. 6a, Extended Data Fig. 10a-b**). We first confirmed that the internal representation of the animal’s current HD during WAKE can be decoded in an unsupervised manner from the manifold as the angular coordinate (‘virtual HD’) of each HD cell population vector on the ring. The HD tuning curves of both PoSub-HD and PoSub-FS cells computed using virtual HD values during WAKE were equivalent to those computed using real HD values (**Fig. 6b, Extended Data Fig. 10c**). During REM sleep the HD system disengages from the outside world while at the same time representing an internally-generated, drifting virtual HD ^11,48^. We thus applied Isomap to REM PoSub-HD population vectors and computed the corresponding virtual HD (**Fig. 6c, Extended Data Fig. 10d-e**). As expected, HD tuning curves of PoSub-HD cells generated internally from the animal’s virtual HD during REM were similar to their WAKE counterparts (**Fig. 6d, Extended Data Fig. 10f**). Importantly, HD tuning of PoSub-FS cells could also be recovered during REM based solely on the virtual HD obtained from the HD ring attractor manifold (**Fig. 6d)**. Taken together, these results indicate that the activity of PoSub-FS cells is restricted by the topology of the HD cell attractor and is largely independent of external inputs.

**Fig 6.**
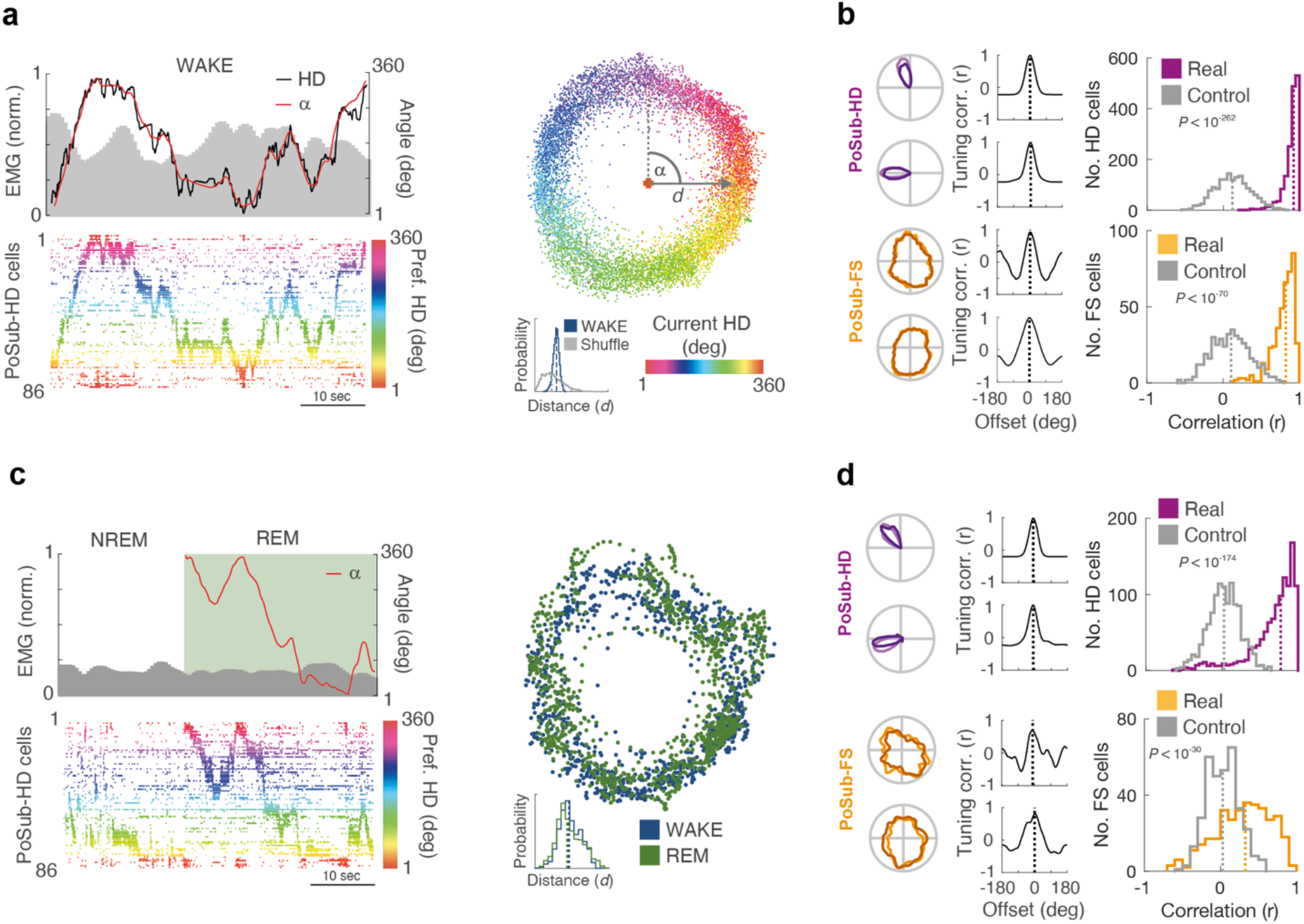
PoSub-FS cells are coupled to the local HD ring attractor manifold during both WAKE and REM. **(a)** Isomap projection of HD cell population vectors during WAKE enables unsupervised reconstruction of HD from the ring manifold. Bottom left: HD cell raster plot of a 50 sec-long fragment of WAKE epoch from a single recording session. HD cells are sorted and color-coded according to their preferred direction. Right: Isomap projection of HD population vectors during WAKE. Each point represents a single population vector, color-coded according to the animal’s current HD. Coordinate *α* is the angular position of each population vector on the ring manifold. Distance of population vectors to the center of the manifold (*d*) is indicative of the ring topology. Top left: Coordinate *α* (virtual HD) precisely matches the animal’s current HD. **(b)** Left: representative examples of PoSub-HD and PoSub-FS cell tuning curve reconstruction during WAKE using the coordinate *α*, and corresponding cross-correlograms between real HD tuning curves (light shades) and WAKE Isomap tuning curves (dark shades). Dotted line shows the offset of maximum correlation. Right: Histograms of the correlation values between WAKE Isomap tuning curves and real HD tuning curves for PoSub-HD cells (top; n = 1602; Wilcoxon signed rank test vs. time-reversed control, Z = 34.7, *P* < 10^−272^) and PoSub-FS cells (bottom; n = 427; Wilcoxon signed rank test vs. time-reversed control, Z = 17.9, *P* < 10^−70^). **(c)** Isomap projection of HD cell population vectors during sleep from the same recording session. Green shading, REM epoch. Bottom left: HD cell raster plot of a 50 sec-long fragment of a sleep epoch from a single recording session. HD cells are sorted and colour-coded according to their preferred direction during WAKE. Right: Isomap projection of HD population vectors during REM and WAKE (sub-sampled). Each point represents a single population vector. Distance of population vectors to the center of the manifold (*d*) is the same during WAKE and REM. Top left: Coordinate *α* (virtual HD) represents the internal representation of direction during REM. Non-REM sleep (NREM) epochs were not analyzed. **(d)** Left: representative examples of PoSub-HD and PoSub-FS cell tuning curve reconstruction during REM using the coordinate *α*, and corresponding cross-correlograms between Isomap tuning curves during WAKE (light shades) and REM (dark shades). Dotted line shows the offset of maximum correlation. Right: Histograms of the correlation values between WAKE Isomap tuning curves and real REM Isomap tuning curves for PoSub-HD cells (top; n = 1148; Wilcoxon signed rank test vs. time-reversed control, Z = 28.2, *P* < 10^−174^) and PoSub-FS cells (bottom; n = 317; Wilcoxon signed rank test vs. time-reversed control, Z = 10.8, *P* < 10^−30^).

## Discussion

In summary, our results establish that PoSub-FS cells, despite non-canonical and symmetrical appearance of their HD tuning curves, share many tuning properties with canonical HD cells: their tuning is stable over time and across environments and is anchored to distal landmarks. However, unlike their canonical counterparts, PoSub-FS cells represent HD in a reciprocal encoding space – the Fourier domain. When this is taken into account, the tuning curves of inhibitory and excitatory cell populations in the PoSub share the same tuning resolution. Finally, we found that this relationship is a local property of the network as the tuning of PoSub-FS cells does not depend on the upstream thalamic input from the ADN and is tightly coupled to the intrinsic dynamics of PoSub-HD cells. We predict that this functional relationship between excitatory and inhibitory cell populations is a general feature of cortical neuronal systems.

FS cells integrate and reflect the activity of anatomically proximal excitatory neurons ^24,27,35,51,52^. This raises the possibility that PoSub-HD neurons are organized in local assemblies representing spatial symmetries. Previous studies have reported symmetries in HD tuning in the retrosplenial cortex ^53^ and in spatial tuning in the medial entorhinal cortex in the form of grid cells ^3^, border ^54^ and band cells ^55^, as well as neurons modulated by environment boundaries ^56^ and axis of travel ^57^ in the subiculum. The retrosplenial cortex, medial entorhinal cortex, and subiculum are three of the main output structures of the PoSub. While HD cell activity in the ADN-PoSub network is crucial for grid cell activity in the medial entorhinal cortex ^58^, it remains to be shown whether the organization of PoSub-HD cells into functional and symmetrical assemblies influences downstream spatial symmetries.

The relationship between excitatory and inhibitory tuning that we observed in the cortical head-direction system may constitute a general principle extending to other cortical systems. Thus, in the primary visual cortex, orientation- and direction-selectivity tuning of excitatory and inhibitory cell populations may show similar equivalence in the Fourier space. What follows is that inhibitory cell tuning may display the same type of symmetries observed here but for orientation and direction. Similarly, we predict that in the medial entorhinal cortex the Fourier signature of grid cell tuning should match the average Fourier signature of the FS cell population within the same module of similarly-spaced grid cells. Moreover, FS cells could be tuned to the spatial frequencies of the underlying toroidal topology of grid cell population activity ^59^.

## Methods

### Subjects

All procedures were approved by the Animal Care Committee of the Montreal Neurological Institute at McGill University in accordance with Canadian Council on Animal Care guidelines. The subjects were adult (> 8 week old) mice bred by crossing wild-type females on C57BL/6J background (Jackson laboratories 000664) with either homozygous male VGAT-IRES-Cre mice (Jackson laboratories 028862, n = 36) or PV-IRES-Cre mice (Jackson laboratories 017320, n = 3). An addition mouse implanted with a Neuropixel probe (**Fig. 1a-c**) was a cross-bred C57BL/6J and FVB (Jackson laboratory 001800). Mice were kept on a 12 hour light/dark cycle and were housed in group cages (2 – 5 mice per cage) before electrode implantation surgery and individually afterwards.

### Electrode implantation

C57BL/6 mice (Jackson Laboratory) were implanted under isofluorane anaesthesia, as previously described ^11^. Briefly, silicon probes were mounted on in-house build movable microdrives and implanted through a small craniotomy. Probes were implanted either vertically above left PoSub (from Bregma: AP, -4.24 mm; ML, 2.05 mm; DV, -1.00 mm) or at a 26 degree angle pointing away from the midline into left posterior retrosplenial cortex (from Bregma: AP, -4.24 mm; ML, 1.70 mm; DV, -1.00 mm). A mesh wire cap was then build around the implanted microdrive and was reinforced with UV-cured adhesive. Mice were allowed to recover for at least 3 days before electrophysiological recordings.

The probes consisted of either a Neuropixel 1.0 probe (384 active sites arranged in a dense checkerboard layout, i.e. two columns, 20 μm between each row), a single shank with 64 recording sites (H5, Cambridge Neurotech, electrodes staggered in two rows, 25 μm vertical separation) or 4 shanks with 8 recording sites each (Buzsaki32, Neuronexus, electrodes staggered in two rows, 20 μm vertical separation). In all experiments both ground and reference wires were soldered to a single 100 μm silver wire which was then implanted 0.5 mm into the cerebellum.

### Recording procedures

During the recording sessions, neurophysiological signals were acquired continuously at 20 kHz on a 256 channel RHD USB interface board (Intan Technologies). The wide-band signal was downsampled to 1.25 kHz and used as the LFP signal. The animals were tethered to a motorized electrical rotary joint (AERJ, Doric Lenses) in order to enable free movement around the enclosure.

Ahead of the main recording session, the microdrive was lowered over several hours in small (35 - 70 μm) increments until the whole shank was positioned in PoSub or ADN. A short open field session was then recorded in order to map the HD receptive fields of all neurons. For PoSub, the recording depth was adjusted so that sharply-tuned HD cells (a hallmark of PoSub) are present along the whole length of the shank. Data collection did not commence until at least 2 hours after the last depth adjustment. Only one recording session per mouse was included in the analysis in order to prevent double counting of cells.

Animal position and orientation was tracked in 3D using 7 infrared cameras (Flex 13, Optitrack) placed above the enclosure and coupled to the Optitrack 2.0 motion capture system. Five small tracking markers were attached to the headcap and additional 2 larger markers were attached to the amplifier chip. In addition, video recording was captured by an overhead camera (Flex 13, Optitrack) placed close to the rotary joint. Animal position and head orientation was sampled at 100 Hz and was synchronized with the electrophysiological recording via TTL pulses registered by the RHD USB interface board (Intan). The tracking system was calibrated in the same manner across all recording sessions to ensure that the tracking coordinates were the same across the whole dataset.

### Behavioural procedures

Before the implant surgery, mice were habituated over several days to forage for small pieces of Honey Cheerios cereal (Nestle) in the open field. For most recordings, the recording chamber consisted of a metal frame (90 × 90 × 180 cm) supporting a plastic platform with removable walls (width: 80 cm, height: 50 cm) that could be arranged into either a square or triangular open field. The recording protocol consisted of a sleep session in the home cage, followed by open field exploration in a square arena and another sleep session. A subset of animals then explored a triangular arena. A white rectangular cue card on one of the walls served as a salient cue. Both environments were oriented so that the wall with the cue card always faced the same direction.

### Optogenetic experiments

Mice were injected with an adeno-associated virus vector (AAV 2/9 CAG-Flex-ArchT-EGFP or CAG-Flex-EGFP, titer: 4-5 × 10^12^ GC/ml, Neurophotonics) into the thalamic reticular nucleus (from Bregma: AP, -0.70 mm; ML, 1.25 cm; DV, -3.25 cm) under isofluorane anesthesia either unilaterally for ADN recordings or bilaterally for PoSub recordings. Injections (300-400 nl per injection site) were done with a microinjector (Harvard Apparatus and Nanofil syringe) through a small craniotomy, at the speed of 100 nl/s. The needle was left in place for 2-5 min after injection in order to minimize backflow.

After at least 3 weeks since the injection surgery, optic fiber implants (Doric Lenses, MFC_200/240-0.22_25mm_SM3) were implanted unilaterally (left hemisphere, ADN recordings) or bilaterally (PoSub recordings) above ADN at a 20-degree angle from the sagittal plane (from Bregma: AP, -0.82 mm; ML, 1.00 cm; DV, -2.25 cm). Mice were then implanted with a microdrive-mounted Buzsaki32 probe above left ADN (Bregma: AP, -0.82 mm; ML, 0.85 cm; DV, -2.00 cm) or either Buzsaki32 or H5 probe above left PoSub, as described above.

Laser light was delivered from a 520 nm fiber-coupled laser diode module (Doric Lenses) controlled with a laser diode module driver (Doric Lenses). Light power output at the tip of the fiber implant was measured before each implantation and a dose-response curve was calculated individually for each implant. Light output was then set to 14-16 mW before each recording session. For this subset of mice, the second sleep session was followed by a second exploration session in the open field, during which a laser stimulation protocol was delivered via patch cords attached to the optic fiber implants. After 5 min of exploration, light pulses (1 s) were delivered at 0.2 Hz in groups of 60 (5 min total), each followed by 5 min of no stimulation. Four such epochs were delivered in total, resulting in 240 light pulses over a 45 min recording session. Such short and interspersed pulses were chosen to promote even sampling of the animal’s current orientation and minimize overheating of the brain.

For animals implanted with single-shank linear probes, only one session per mouse was included in the analysis in order to prevent double-counting of cells. For animals implanted with 4-shank Buz32 probes, multiple sessions per mouse (obtained on separate days) were included in the analysis, ensuring that the probe was moved by at least 70 μm between the recording sessions.

### Cue rotation experiment

A subset of animals implanted into PoSub with linear H5 probes underwent a cue rotation experiment in a separate recording session. To this end, the frame of the recording chamber was fitted with black plastic insets that covered the floor (90 × 90 cm) and walls in order to hide any visual cues. Additionally, the outside of the recording chamber was covered with thick fabric in order to block off any light coming from the outside. Each of the four walls had an identical panel made of two LED strips (yellow V-shape or blue X-shape) in the centre. A small (30 cm diameter) elevated circular platform was placed in the centre of the arena, at the elevation corresponding to the bottom of the LED panels. Before each recording session, two adjacent LED panels were chosen as distal visual cues. The LED light intensity was chosen so that the panels are visible in the dark but provide minimal illumination of the surrounding area. The on/off cycle of the LED panels was controlled with an Arduino computer. In order to eliminate other sensory cues, the whole recording chamber was thoroughly cleaned with antibacterial wipes before the experiment and white noise was emitted from speakers placed underneath the chamber, below the circular platform.

The recording protocol consisted of a 1-hour sleep session in the home cage placed on the circular platform, followed by a 75 min cue rotation session. At the start of the cue rotation session a single LED cue dimly illuminated, the mouse was placed directly on the circular platform and was left undisturbed for the duration of the session. The recording chamber was then sealed to ensure near-complete darkness except for the dimly illuminated LED cue. After 15-20 min of exploration with a stable cue, the cue rotation protocol was initiated. The protocol consisted of the illuminated cue switching 8 times back and forth between two adjacent walls (16 total cue rotations), each time staying in place for 200s. Additionally, in order to habituate the mouse to the cue disappearing from its field of view, the cue was turned off for 0.1 s every 20 s. In order to encourage the mouse to explore the platform for the whole duration of the session, the experiment was conducted in the middle of the dark phase of the light cycle and the recording chamber was sprayed with a novel odour (scented air freshener) right before the cue rotation session.

### Tissue processing and probe recovery

Following the termination of the experiments, animals were deeply anesthetized and perfused transcardially first with 0.9% saline solution followed by 4% paraformaldehyde solution. The microdrive was then advanced to remove the probe from the brain and the probe was moisturized with distilled water while the brain was being extracted. Brains were sectioned with a vibratome sagitally in 40 μm thick slices. Sections were washed, counterstained with DAPI and Red Neurotrace and mounted on glass slides with ProlongGold fluorescence antifade medium. Sections containing probe tracts were additionally stained with a Cy3 anti-Mouse secondary antibody to help visualize the electrode tract. Widefield fluorescence microscope (Leica) was used to obtain images of sections and verify the tracks of silicon probe shanks, optic fiber position and virus expression.

Probes were lifted out of the brain immediately after perfusion by turning the Microdrive screw all the way up. As the headcap was being manually dismantled, the probe shank was then kept from drying by infusions of distilled water into the headcap. Once the drive-mounted probe was separated from the headcap, it was immersed in 3% peroxide for 5 minutes and rinsed with distilled water. The probe was then dipped in and out of a warm Contrad solution for several minutes, followed by a 2-hour incubation in a warm 2% Tergazyme solution. The drive-mounted probe was then rinsed with distilled water and stored for the next implantation. This multistep cleaning protocol enabled us to implant an individual probe an average of 3 times.

### Spike sorting and unit classification

Spike sorting was performed semi-automatically, using Kilosort 2.0 ^60^ followed by manual curation of the waveform clusters using the softwares Klusters ^61^ and Phy ^62^. At this stage, any cluster without a clear waveform and clear refractory period in the spike train autocorrelogram were classified as noise and cluster pairs with similar waveforms and common refractory period in their spike train cross-correlogram were merged.

For PoSub recordings, viable units were first identified as units that (1) had an average firing rate of at least 0.5 Hz during open field exploration, and (2) had a waveform with negative deflection (criterion aiming to exclude spikes from fibers of passage). Next, putative excitatory cells and putative FS interneurons were classified on the basis of their mean firing rate during open field exploration and the through-to-peak duration of their average waveforms (**Extended Data Fig. 1f**). Putative FS interneurons were defined as cells with short trough to peak duration (< 0.38 ms) and high mean firing rates (> 10 Hz). Conversely, cells with long trough-to-peak (>0.38 ms) and low mean firing rates (< 10 Hz) were classified as putative excitatory cells.

### HD tuning curves and tuning metrics

The animal’s HD was calculated as the horizontal orientation of a polygon constructed in Optitrack from the 3-dimensional coordinates of all tracking markers, relative to the global coordinates, constant across the whole study (see above). In order to minimize the effect of animal’s velocity on HD tuning curves, the dataset was limited to the epochs when animal’s speed exceeded 2 cm/s for all analyses except cue rotation and optogenetic experiments, where epoch duration (200 and 240 s, respectively) was too short to warrant further refinement. HD tuning curves were then computed as the ratio between histograms of spike count and total time spent in each direction in bins of 1 degree and smoothed with a Gaussian kernel of 3° s.d.

Since a sizable proportion of HD cells in our dataset had tuning curves with multiple peaks and therefore low resultant vector lengths, we chose to define HD cells based on a HD information ^63^, calculated for N angular bins as: 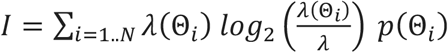 where *λ*(Θ_*i*_) is the firing rate of the cell for the *i*th angular bin, *λ* is the average firing rate of the neuron during exploration and *p*(Θ_*i*_) is the occupancy (i.e. normalized time spent) in direction Θ_*i*_.

For each cell, we obtained the control tuning curve by computing a tuning curve using time-reversed HD angle – a method that preserves the dynamics of both the spike train and the HD angle but decouples the two from each other. We then classified HD cells as those with HD information scores higher than 99^th^ percentile of the null distribution (> 0.2 bits/spike, 85% of putative excitatory cells).

Similarly, we used HD information (see above) to characterize the HD tuning of FS cells. Should we apply our HD cell definition to the recorded putative FS cells, 67% of FS cells would pass the 99^th^ percentile of the null distribution (> 0.015 bit/spike). However, since the tuning of the vast majority of recorded FS cells showed positive correlation across two halves of the recording session (**Fig. 1g**), we decided to include all recorded FS cells in the subsequent analyses.

### Cross-validated HD tuning curve auto- and crosscorrelograms

We obtained HD tuning curve auto- and crosscorrelograms by computing Pearson’s correlation coefficients between the reference tuning curve vector and the second tuning curve vector (from either the same or another cell), which was circularly shifted by 0 to 359 bins. In order to minimize the effect of non-HD factors on tuning curves computed from the same epoch, we employed a cross-validation procedure whereby the two tuning curves were computed from separate halves of the epoch.

When computing the HD tuning curves during exploration of the triangular open field, we have noticed that sometimes the cells’ receptive fields were rotated with respect to the prior square open field exploration. In order to counteract this, we first calculated for each HD cell the degree of tuning curve rotation between the two environments via cross-correlation. We then used the average rotation value per recording session to circularly shift all triangular open field tuning curves in the opposite direction by the equivalent amount. This allowed us to compute the true tuning curve correlation between the two environments.

### Detection of mono-synaptic connections

Mono-synaptic connections between PoSub-HD and PoSub-FS cells were detected as described previously (Peyrache et al, 2015). Briefly, spike train cross-correlograms of +/- 50 ms binned in 0.2-ms windows were convolved with a Gaussian kernel of 4 ms s.d, resulting in a predictor of the baseline rate. At each time bin, the 99.9 percentile of the cumulative Poisson distribution (at the predicted rate) was used at the statistical threshold for significant detection of outliers from baseline. A putative connection was considered significant when at least two consecutive bins in the cross-correlograms passed the statistical test.

### Analysis of HD cell realignment after cue rotation

In order to estimate the degree of realignment of the HD system following cue rotation, the ‘internal’ HD was reconstructed using a Bayesian framework following previous work ^11,64^. In short, HD cell spike times were binned into population vectors (50 ms window, smoothened in 100 ms s.d. Gaussian windows). Based on cells’ tuning curves from the period of exploration with stable cue, the population vectors were converted into a probabilistic map under the hypothesis that HD cells fire as a Poisson process. The instantaneous internal HDs were taken as the maxima of these probabilistic maps. These estimates faithfully tracked the head orientation of the mouse during the period preceding cue rotation. The degree of realignment of the HD system was calculated as the decoder error: angular difference between the real HD and the internal HD at each time bin. Since not all cue rotations resulted in HD realignment, we excluded all cue rotation epochs that (1) resulted in less than 45 degrees of mean decoder error in the following 200 s epoch, and (2) occurred when the animal was stationary in the preceding epoch (average velocity < 2 cm / sec).

We have observed that in our paradigm the realignment was not instantaneous but often lasted for several seconds, independently of the degree of Gaussian smoothing applied to the spike trains. In order to estimate the rate at which the HD system remaps, we fitted a sigmoid curve to the decoder error values following each cue rotation using the *sigm_fit* function (https://www.mathworks.com/matlabcentral/fileexchange/42641-sigm_fit). We defined the beginning and end of realignment epochs as the timestamps corresponding to the values of 0.01 and 0.99 of the normalized sigmoids. We then, for each cell, calculated a HD tuning curve for the remainder of each cue epoch (from the end of realignment to next cue rotation), and computed the cross-correlation (see above) between HD tuning curves from consecutive epochs. For each cell, the degree of realignment was defined as the tuning curve offset which results in highest correlation coefficient. The difference in realignment between FS and HD cells was defined as, for each FS cell, the angular difference between its degree of realignment and the average realignment of HD cells in the same epoch.

### Classification of sleep states

Sleep scoring was performed using the automated SleepScoreMaster algorithm ^65,66^ (Buzsaki laboratory, https://github.com/buzsakilab/buzcode/tree/master/detectors/detectStates/SleepScoreMaster). Briefly, the wide-band signal was downsampled to 1.25 kHz and used as the LFP signal. Electromyograph (EMG) was computed from correlated high-frequency noise across several channels of the linear probe.

### Pairwise spike rate coupling

Quantification of pairwise spike rate coupling between cells was quantified using a General Linear Model (GLM) according to the method described in ^49^. Spike trains were binned in 100 ms bins and smoothened in 100 ms s.d. Gaussian windows. The population firing rate was calculated by aggregating all spike times from all recorded units in a given recording and processing them in the same manner as single spike trains. Both binned trains were then restricted to either WAKE or REM epoch (see above).

The GLM was fitted using the MATLAB ‘glmfit’ function. The binned spike train of cell A was modelled as a Poisson process, as a function of both the binned spike train of cell B and the binned population firing rate, using a log link function. The model produced a coefficient of coupling between the spike trains of cells A and B (‘Beta’), as well as a coefficient for the coupling of cell A to the population firing rate (‘Beta-pop’). The procedure was repeated by offseting the spike train of cell A by ± 10 seconds in 100 ms intervals, to yield the equivalent of spike train cross-correlogram that discounts the coupling of cell A to the local population rate. Since this procedure, unlike Pearson’s correlation, is not symmetric, it was repeated by substituting cell A and cell B and averaging the coupling coefficient values at equivalent offset intervals. A cell pair was removed from the analysis in rare cases when the ‘glmfit’ function identified the model as ill-fitted or reached the iteration limit.

### Visualization and analysis of the ring manifold

For visualizing the HD manifold during WAKE, HD cell spike times from the whole epoch were binned into population vectors in 200 ms bins, smoothened in 400 ms s.d. Gaussian windows and a square root of each value was taken. Then, non-linear dimensionality reduction was performed using the Isometric Feature Mapping (Isomap) algorithm ^39^ implemented in the Matlab Toolbox for Dimensionality Reduction (https://lvdmaaten.github.io/drtoolbox/). The parameters were set to 12 nearest neighbours and 3 dimensions – the latter to inspect if there is no higher dimensional manifold in the data. Shuffled Isomap embeddings were computed by shifting each cell’s binned spike train in time by a random number of bins.

Internal HD at each time bin was then calculated as a four-quadrant arctangent of the two Isomap dimensions (range: -180° to 180°). Importantly, the internal HD generated this way has arbitrary directionality (clockwise/anticlockwise) and an arbitrary point of origin. To calibrate it, Isomap directionality was established by computing the Isomap error as the difference between real HD values and (1) internal HD values and (2) internal HD multiplied by -1, and selecting the directionality with error of smaller circular variance. Internal HD tuning curves were then computed as the ratio between histograms of spike count and total time spent in each internal HD in bins of 1 degree and smoothed with a Gaussian kernel of 3° s.d. Real HD tuning curves were computed by downsampling the real HD into 200ms bins and applying the same procedure as above. In order to correct for the arbitrary point of origin, HD tuning curve cross-correlations (see above) were computed between each real and internal HD tuning curve of each HD cell, and the mean offset of maximum correlation was then used to circularly shift all tuning curve vectors by the equivalent number of bins. While this procedure (as well as Isomap mapping in general) was dependent on the real HD tuning of HD cells, it was independent of FS cell tuning. Control HD tuning curves were computed by time-reversing the Isomap angle (see above).

For the comparison between internal HD tuning curves during WAKE and REM, population vectors across WAKE and REM were computed in the same manner as above. WAKE population vectors were then randomly downsampled to equal in number to the REM population vectors. Isomap algorithm was then run on both WAKE and REM population vectors together. Internal HD was computed in the same way as above. HD tuning curves were computed in bins of 6 degrees and smoothed with a Gaussian kernel of 1° s.d for both real and internal HD. Larger bin size was chosen due to sometimes uneven sampling of virtual HDs during REM.

### Fourier analysis of HD tuning curves

To decompose tuning curves into Fourier series, we projected the tuning curves onto a basis of sine and cosine functions whose frequencies were the harmonic of the unit circle, i.e. from the fundamental frequency (period of 360°) to the highest possible frequency (2°, the inverse of the Nyquist frequency as tuning curves are computed in 1° bins). The power, or Fourier coefficients, at a particular frequency was defined as the root mean square of the projection values onto the sine and cosine basis at that frequency. Similarly, the phase was defined as the arctangent of the projections. The validity of the projection was verified by checking that the sum of squared Fourier coefficients is equal to the variance of the tuning curves (Parseval’s identity), which was indeed the case (**Extended Data Fig. 3e**). Since higher Fourier components likely represent noise fluctuations in the tuning curves, we focused our analysis on the relative power of the first ten Fourier components, normalizing their individual power values to the sum of their power.

Kullback–Leibler (KL) divergence was used as a measure to assess the similarity between the individual Fourier spectra and the population means. While KL divergence is regularly used to compare probability distributions, we deemed it appropriate to apply it to normalized Fourier spectra as they were mathematically indistinguishable from probability distributions. We thus computed the KL divergence between the spectrum of an individual neuron *σ*_*i*_(*k*) (with *k ∈* [1. .10] the angular frequencies) and the average Fourier spectrum of a population *S*_*i*_(*k*) as follow:

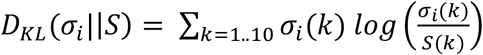

While KL divergence is not symmetrical, i.e. *D*_*KL*_(*σ*_*i*_||*S*) does not equal *D*_*KL*_(*S*||*σ*_*i*_), we always apply it in the same direction, i.e. *D*_*KL*_(*σ*_*i*_||*S*).

### Isomap analysis of HD tuning curve auto-correlograms

Cross-validated autocorrelograms of each cell’s tuning curve were computed as described above. Then, non-linear dimensionality reduction was performed using the Isometric Feature Mapping (Isomap) algorithm ^39^. The parameters were set to 12 nearest neighbours and 2 dimensions. When mapping the first three Fourier components onto the resulting embedding, we normalized their power to the total sum of their powers.

### Anatomical tract tracing

VGAT-Cre mice were injected with an adeno-associated virus vector (AAV 2/9 CAG-Flex-EGFP, titer: 4-5 × 10^12^ GC/ml, Neurophotonics, 500 μl per injection site) bilaterally into the thalamic reticular nucleus as described above. Four weeks after injections, animals were perfused transcradially with 4% PFA in phosphate-buffered saline and their brains were then cut coronally in 40 μm sections with a freezing microtome. The sections were counterstained with blue NeuroTrace and mounted on sides with Coronal z-stacks of the sections containing the rostral thalamus were taken with a Leica SP-8 confocal microscope at x10 magnification, using the same settings for all sections. GFP signal was acquired using the 473 nm excitation laser line. Z-projections of each stack were then obtained using ImageJ software. Quantification of anterograde tracing was done in ImageJ. The images were converted to grayscale and rectangular regions of interest (ROI) were defined within each thalamic nucleus. Average pixel intensity per ROI was then calculated using the ‘Measure’ function.

### Gain modulation analysis

Epochs of optogenetic stimulation (light ON) consisted of time periods when the laser was switched on (240 pulses of 2 sec duration, 240 sec per session). Control epochs (light OFF) were defined as time-periods in-between light pulses (240 periods of 4 sec duration, 960 sec per session). Light ON and light OFF tuning curves were computed from these periods. For analysis of tuning curve width, HD tuning curves were then computed in bins of 1 degree and smoothed with a Gaussian kernel of 3° s.d. For analysis of additive and multiplicative gain, in order to preserve the independence of individual angular bins, HD tuning curves were then computed in bins of 6 degrees with no Gaussian smoothing applied. Light ON and light OFF tuning curves for each cell were then normalized by dividing them by the maximum value of the light OFF tuning curve. To calculate the additive and multiplicative factors for each cell, a linear fit between light ON and light OFF tuning curve vectors was then obtained using the Matlab *polyfit* function. The slope of the resulting linear fit and its Y intercept were then taken as multiplicative and additive gain values, respectively.

### Data analysis and statistics

All analyses were conducted using software custom-written in Matlab R2020b (Mathworks). Unless otherwise specified, statistical comparisons were performed with Mann-Whitney U test, Willcoxon Signed Rank Test or ANOVA with multiple comparisons, where applicable. All statistical tests are two-tailed.

### Simulations - methods

The linear model linearly sums input tuning curves with randomly distributed weights. We used the recorded smoothed excitatory cells tuning curves as inputs (concatenating ADn and PoSub). Similar results could be obtained using Normal, Uniform or Log-Normal weights distribution. We also verified that sparsifying these weights did not change the simulation results. Sparsification was applied by setting to 0 a subset of the weights after random sampling.

### Simulations - theory

We theoretically derive the asymptotic behavior of the proportion of simulated cells with a certain Fourier power. We show that this proportion becomes independent of the weights mean and standard deviation in the limit of low or high variance relative to the mean. This proportion is a statistic of the ratios of each Fourier power over the sum of all Fourier power. Consequently, it is sufficient to show that this ratio converges to a limit distribution independent of the weights mean and variance.

Here, we sought to determine the average response of neurons to an encoded feature *θ* (i.e. its tuning) as a function of the tuning of its inputs and their synaptic weights. At first, we assume that the response of the neuron *y*_*j*_ is linear and free of noise. In this case, the tuning curve is

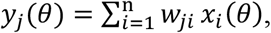

where *x* are the tuning curves of presynaptic neurons, *θ* is the feature, *w*_*ji*_ the synaptic weight from input neuron i to output neuron j, and *n* the number of input neurons.

Let us rewrite each weight as *w*_*ji*_ = *μ*_*w*_ + *v* _*ji*_*σ*_*w*_

Where *v*_*ji*_ is the z-scored weight 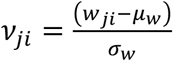

For a set of n input tuning curves *x*(*θ*),

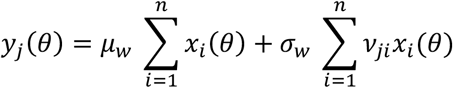

For normally distributed weights, the distribution of *ν*_*ji*_ is independent of the mean and standard variation of the weight distribution. Consequently 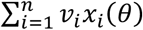 is independent of the mean and standard deviation. Let us write the Fourier coefficients of this term as *b*_*l*_, so that we have:

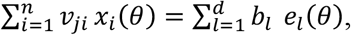

where e is the discrete Fourier basis:

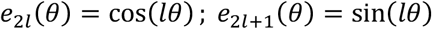

Noting the Fourier coefficients of the sum of input tuning curves as *a*_*l*_ :

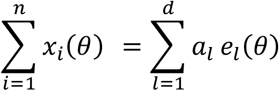

the Fourier coefficients of the output tuning curves can be expressed as:

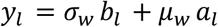

Finally, the Fourier power of the output tuning curves are:

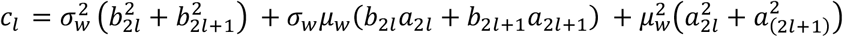

In the limit *σ*_*w*_ ≫ *μ*_*w*_ *or σ*_*w*_ ≪ *μ*_*w*_, the ratio of Fourier power *r*_*l*_ will only depend on the distribution of the *b*_*l*_ or of the *a*_*l*_.

If *σ*_*w*_ ≫ *μ*_*w*_, we have 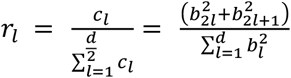

And if *σ*_*w*_ ≪ *μ*_*w*_ we have 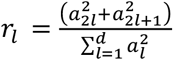

For normally distributed weights, *b*_*l*_ and *a*_*l*_ are independent of the weight mean and standard deviation (by definition). Therefore, the ratio of Fourier power converges to a distribution that is independent of the weight mean and standard deviation. By extension, the proportion of cells in the data showing maximum power in a certain fold is independent of the weight statistics. In the case *σ*_*w*_ ≫ *μ*_*w*_, the power ratio depends only on the spectrum of the weighted sum of input tuning curves.

We next demonstrate that the power ratio of the output tuning curves is entirely determined by the input tuning curve spectrum – independent of the weight mean and standard deviation, as well as the choice of weight distribution.

We show that in the limit of a large number of input cells, the Fourier coefficients of the outputs tuning curves are distributed as independent normal distributions, even for non-normally distributed weights.

#### Distribution of FS cells tuning curve

We asked the question of what the resulting distribution of output neuron tuning curves is in the linear regime.

We first present the case of a linear combination with normally distributed weights *W* (mean *μ*_*W*_ and standard variation *σ*_*W*_) and then generalize it to other weights.

#### Demonstration for normally distributed weights

Tuning curves are discretized in *m*_*θ*_ points and this discretization is as fine as needed to preserve all the information of the tuning curves. The exact distribution of tuning curves can be derived by projecting the input tuning curves in their natural basis i.e by decomposing them with a singular value decomposition (SVD). The (*n, m*_*θ*_) tuning curve matrix X is expressed as *X* = *USV*^*T*^.

To determine the statistics of the output tuning curve *y* = *WX*, one can project *y* onto the singular vectors of *X*. The projection of *y* onto the *l-*th singular vector is:

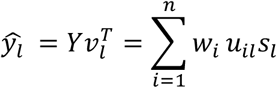

It is sufficient to know the value of the first *d* = *rank*(*X*) projection 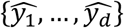 to fully reconstruct the tuning curve in the feature space: 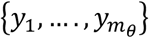. Indeed, the projection onto the *d* + 1, …, *m*_*θ*_ singular vectors are null. Therefore, the distribution of tuning curves can be equivalently described either in feature space or singular component space.

This projection 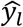 is an affine transformation of the random multivariate normal variable *W* by *US*_(:,1:*d*)_. As *US*_(:,1:*d*)_ forms an orthogonal basis, the 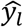 are independent (up to *l* = *d*). As *w*_*k*_ are independent samples from a normal probability distribution, the random variable 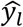 is itself distributed according to a normal distribution 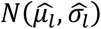 where:

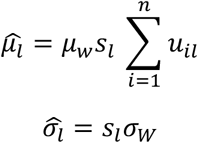

In summary, and quite intuitively, tuning curves of randomly connected linear neurons are constrained to lie in the subspace spanned by the singular components of their input tuning curves. They are distributed in this subspace according to a multivariate normal distribution with diagonal covariance. The weight standard deviation and *l-*th input singular value control the variance of 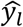.

The above result was derived for normal distributions. For non-normally distributed weights (as it is the case of lognormally distributed positive weights), one potential approach would be to use the central limit theorem (CLT), on the sequence of random variable *w*_1_*u*_1*l*_, …, *w*_*n*_*u*_*nl*_. However, the CLT assumes identically distributed random variables, which is not the case here because the *u*_*il*_ are not identically distributed.

#### Extension of the proof to non-normally distributed weights

Here, we show that the main result holds anyway for non-normally distributed weights in the limit of large number of input excitatory cells n, under some realistic assumptions on the *u*_*il*_. To this end, we use a generalized version of the CLT, the so-called Lyapunov Central limit theorem.

The Lyapunov central limit theorem is the following:

Suppose *V*_1_, … *V*_*n*_ are independent random variable with mean 0 and finite variance 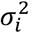.

Let 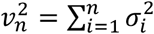 denote the sum of their variance and *S*_*n*_ = *V*_1_ + … + *V*_*n*_.

If for some *δ* > 0 the so-called Lyapunov condition is verified:

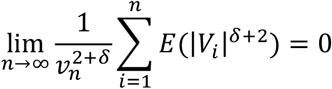

Then 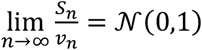 in distribution.

We will apply the theorem to each projection of the output tuning curves onto the right singular vectors of the input tuning curves. This will demonstrate the normality of each projection. We will then prove the independence of these projections.

Let *l* be the index of the right singular vector of the input tuning curves. We note *W*_*i*_ the random variable describing the weights of the *i-*th input.

The dependence of the tuning curve left singular vector on the number of cells can be made explicit:

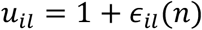

For example, for a set of von Mises tuning curve (i.e., canonical HD tuning curves) with preferred direction uniformly spread on the circle, we have:

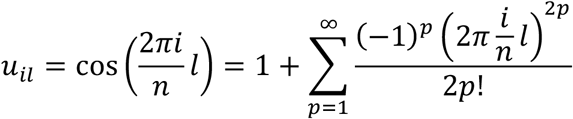

The projection of an output tuning curves on the *l-*th singular vector is:

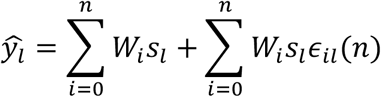

The term 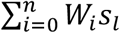 is a sum of identically independently distributed weights and the CLT will apply.

For the second term 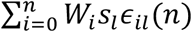 we demonstrate that the Lyapunov condition holds under certain assumption for the *∈*_*il*_(*n*).

We define the random variable *V*_*i*_ = (*W*_*i*_ – *μ*_*w*_)*s*_*l*_*∈*_*il*_(*n*) of mean 0 and finite.

First, we compute the variance of the sum:

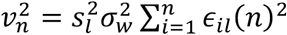

We have

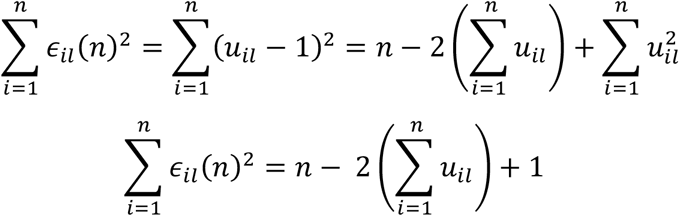

Assumption 1: 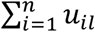 is finite when *n* grows to infinity, i.e 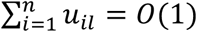

*Note:* this is the case for the set of perfect von Mises tuning curves, as 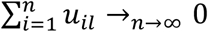

Under assumption 1:

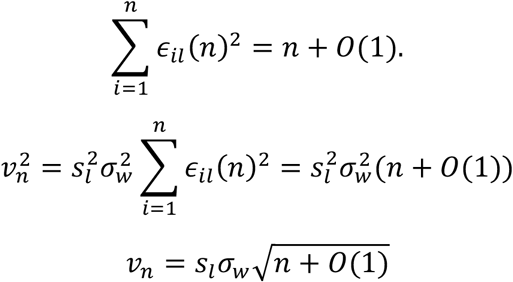

Next let us compute the sum of expectation term. For every *i* ∈ [1, *n*]

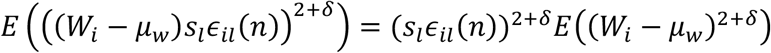

The term *E*((*W*_*i*_ – *μ*_*w*_)^2+*δ*^) is a statistic of the weights, independent of the random variable (i.e., independent of the index *i*).

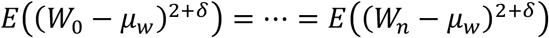

Consequently

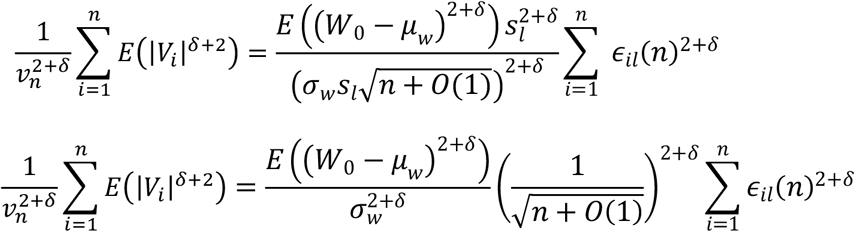

Assumption 2: 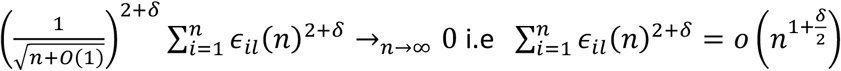

*Note*: Let us show that assumption 2 holds for von Mises tuning curve:

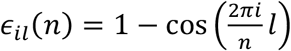

The sum 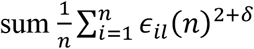 is in that case a Riemann sum that converges to the following integral:

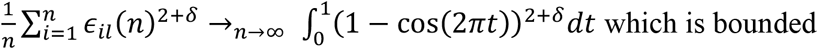

And therefore:

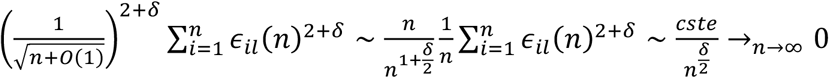

Under the assumption 2, which we proved valid for a theoretical model of perfect input tuning curves, we have consequently:

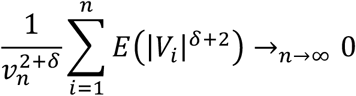

This demonstrates the Lyapunov condition for the *l-*th projection. This proves that 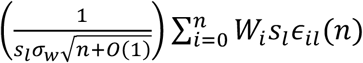 are normally distributed in the limit of large n, and consequently: 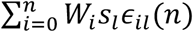 is also normally distributed in the limit of large n.

Therefore we obtained that both 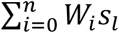 and 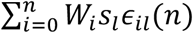 are normally distributed, this proves that 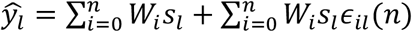 is also normally distributed.

We can compute the mean and variance of this normal distribution:

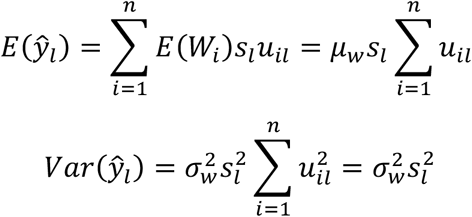

These statistics are true for all values of n, but the distribution itself converges to a normal distribution only at large n. The expectation and the variance of the limit distribution will be the limit of the expectation and variance computed above.

The limit statistics are:

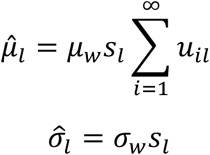

Finally, the independence of the different projections simply stems from the fact that the covariance matrix is diagonal, as this property is the same as for the normally distributed weights. Therefore, 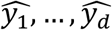 are indeed independent. The output tuning curve projection on the right singular vector of the tuning curves are thus normally distributed and independent.

For canonical tuning curves, modeled as von Mises’ distribution, the singular vectors are exactly the Fourier basis up to a proportionality coefficient. Here, we thus have demonstrated that, under realistic assumptions, the Fourier components of output tuning curves are independent and normally distributed with a variance depending on the weight variance. This is exactly what is seen in the data: while all PoSub-HD cells have qualitatively the same Fourier spectrum, each PoSub-FS cell has a unique spectrum, many of which concentrating power in only one or two components.

## Acknowledgements

We would like to thank Lynda Mainville for technical support. We are thankful to Blake Richards, Mark Brandon, Stuart Trenholm and the members of Peyrache laboratory for comments on the earlier version of the manuscript.

## Author contributions

Conceptualization: AJD, AP; Methodology: AJD, AP, PO; Software: AJD, PO, AP; Validation: AJD, AP; Formal analysis: AJD, AP, PO, EO, EHB; Investigation: AJD, SSC, GV; Resources: APl; Data Curation: AJD; Writing - Original Draft: AJD, AP; Writing - Review & Editing: AJD, AP, PO, ERW; Visualization: AJD, PO; Supervision: AP; Project administration: AP, AJD; Funding acquisition: AP, AJD

## Data and code availability

The datasets generated in this study will be made available online at time of publication. Code will be available upon request.

**Extended Data Fig. 1.**
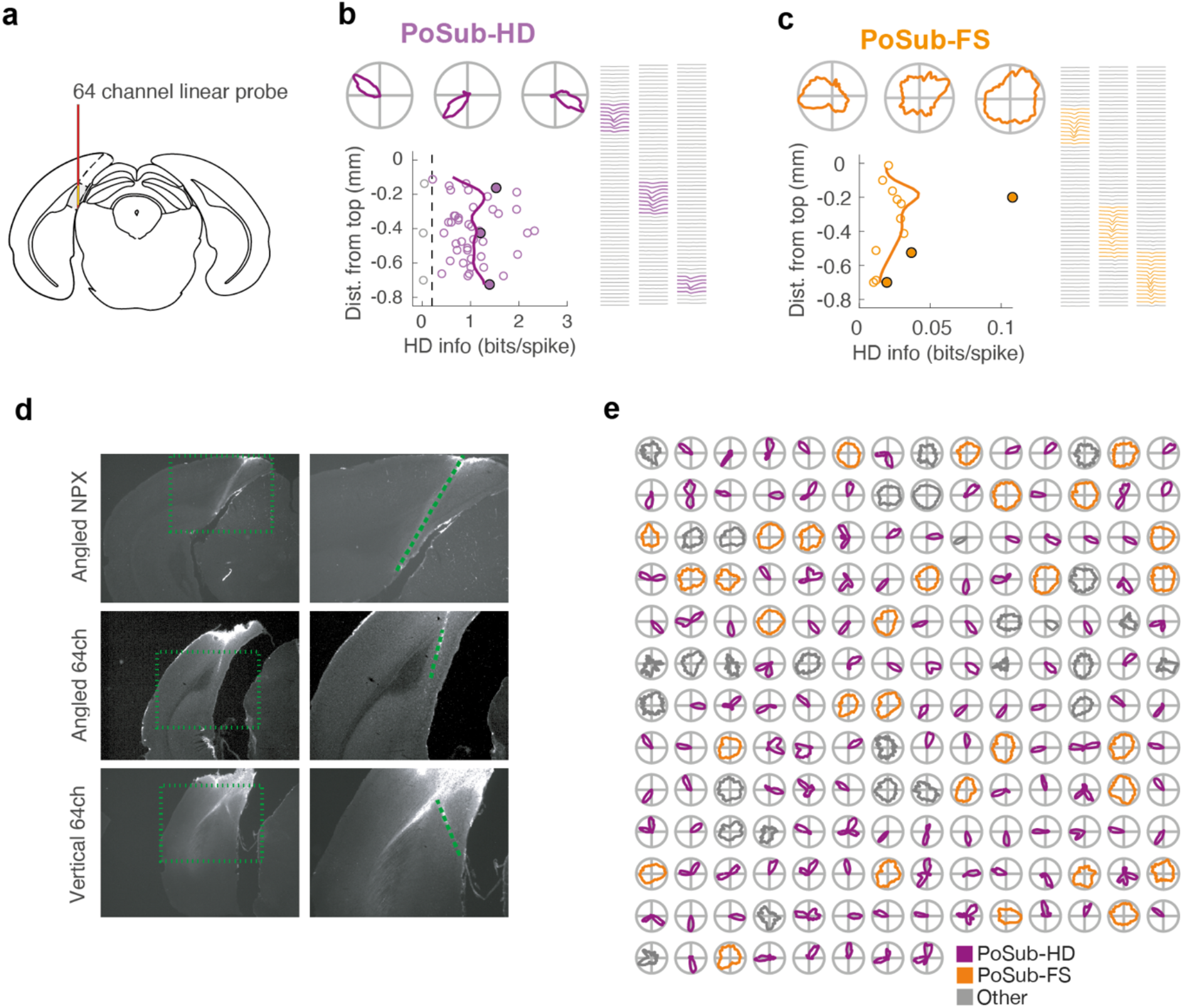
Vertical recordings, examples of probe trajectories and examples of tuning curves. (**a**) Brain diagram showing the vertical positioning of the 64-channel linear probe in PoSub. (**b**-**c**) Scatterplot depicting HD information of all putative excitatory cells (**b**) and putative FS cells (**c**) in a single vertical 64-channel probe recording as well as the running average (solid lines). Representative HD tuning curves and spike waveforms correspond to filled circles. (**d**) Anti-mouse antibody-stained coronal sections depicting the probe tract for the Neuropixel (NPX) recording (top) as well as representative tracts for the angled (middle) and vertical recordings (bottom). Images on the right are magnified regions denoted with a dotted square the left. Dashed line, position of the probe recording sites. (**e**) Tuning curves of all cells recorded in the session depicted in Fig 1D-e, arranged in anatomical order from the top to the bottom of the probe.

**Extended Data Fig. 2.**
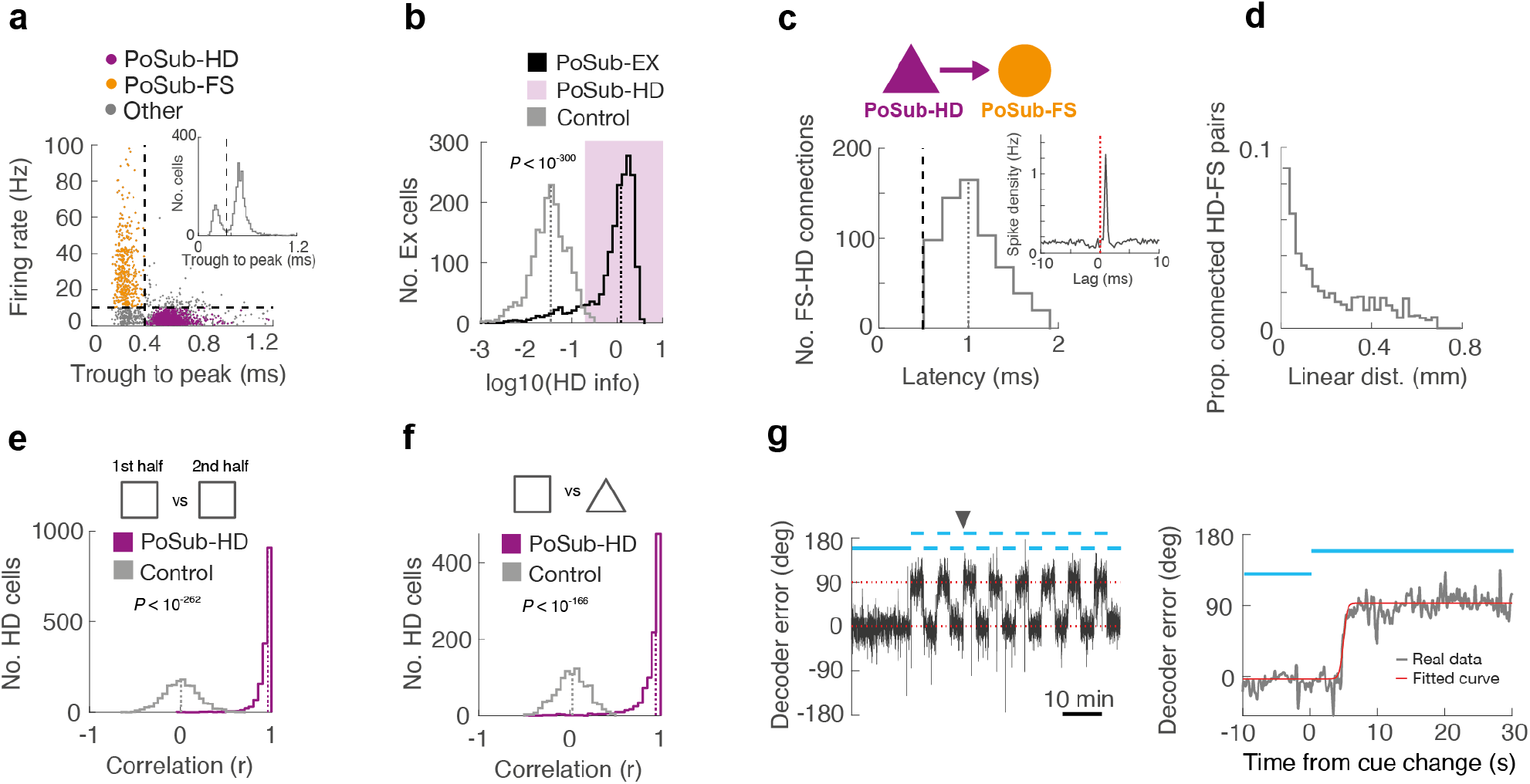
Cell types, synaptic connections, PoSub-HD cell stability and cue rotation. (**a**) Identification of cell types. Left: PoSub units were separated into putative excitatory and PoSub-FS cells based on firing rate and waveform shape (dashed lines). Left, inset: Histogram of trough-to-peak duration in the recorded PoSub unit waveforms. (**b**) Histogram of HD information carried by tuning curves of putative excitatory cells in PoSub (n = 1835; Wilcoxon signed rank test vs time-reversed control, Z = 37.1, *P* < 10^−300^). A sub-population of putative excitatory cells with HD information higher than 99^th^ percentile of the time-reversed control population was classified as PoSub-HD cells (purple shading). Dotted lines show the medians of the depicted distributions. (**c**) Histogram of the latency of putative synaptic connections between PoSub-HD cells and PoSub-FS cells. Dashed line depicts the latency threshold for synaptic connections. Left, inset: example spike-timing cross-correlogram between a PoSub-HD and a PoSub-FS cell showing a putative synaptic connection. (**d**) Histogram depicting probability of an excitatory synaptic connection between PoSub-HD and PoSub-FS cells as a function of linear distance on the electrode array. (**e**) Histogram showing the correlations between PoSub-HD cell tuning curves in two halves of a single recording epoch (n = 1602; Wilcoxon signed rank test vs time-reversed control, Z = 34.7, *P* < 10^−262^). Dotted lines show medians of the depicted distributions. (**f**) Histogram showing the correlations between PoSub-HD cell tuning curves in square and triangle arenas (n = 1013; Wilcoxon signed rank test vs time-reversed control, Z = 27.6, *P* < 10^−166^). Dotted lines show medians of the depicted distributions. (**g**) Angular error of the Bayesian decoder during a representative cue rotation session. Left: rotation of the cue by 90 degrees reliably corresponds to the equivalent change in HD decoding error. Right: changes in decoder error during a single cue rotation epoch (denoted with a black arrowhead on the left). Red line is the sigmoidal function fitted to estimate the end of remapping. Blue bars denote the cue position.

**Extended Data Fig. 3.**
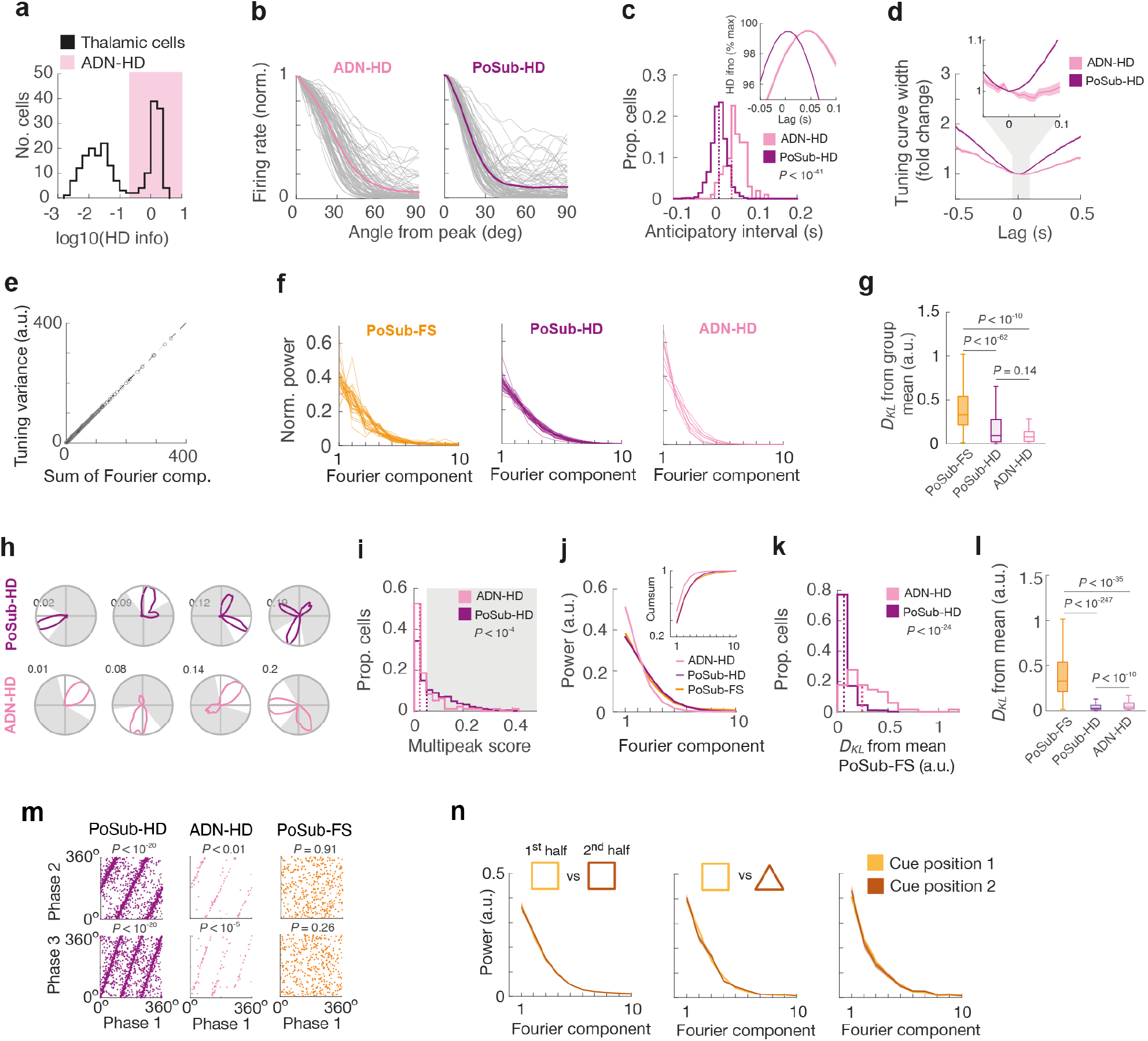
ADN-HD cells, anticipatory intervals and additional analyses of the Fourier signatures. (**a**) Histogram showing bimodal distribution of HD information in the anterior thalamus. Pink background denotes classified ADN-HD cells (n = 97), the threshold of 0.2 bits/spike is the same as that used to classify PoSub-HD cells. (**b**) Tuning curve slopes of ADN-HD cells (n = 97) and PoSub-HD cells (n = 97, randomly selected). Thin gray lines denote individual tuning curves, thick colored lines denote averages of depicted populations. (**c**-**d**) Effect of the anticipatory interval on HD tuning in ADN and PoSub. (**c**) Histogram showing distribution of anticipatory intervals of PoSub-HD and ADN-HD cells (Mann-Whitney U test, n = (1602, 97), Z = 13.6, *P* < 10^−41^). Dotted lines show the medians of the depicted distributions. Inset: change in HD information as a function of time lag. Shaded lines represent mean +/- SEM. (**d**) Tuning width of PoSub-HD and ADN-HD cells as a function of time lag. Shaded lines represent mean +/- SEM. (**e**) Scatter plot showing the correlation between the sum of Fourier components and tuning curve variance. (**f**) Average Fourier spectra of PoSub-FS, PoSub-HD and ADN-HD cells in individual mice. (**g**) Box plots showing statistical distance between individual Fourier spectra and corresponding population averages (ANOVA, cell type, F_(2,2123)_ = 153, P < 10^−60^). Boxes represent the interquartile range (IQR), whiskers represent minimum and maximum values that are not outliers. Outliers were defined as values more than 1.5 x IQR away from the top or bottom of the box. Individual comparisons (Bonferroni correction) displayed on the panel. (**h**-**l**) Influence of multi-peaked HD cells on Fourier signatures of PoSub-HD and ADN-HD cell populations. (**h**) Examples of PoSub-HD and ADN-HD cell tuning curves and their multipeak score. Shaded areas of the polar plots represent the regions outside of the primary receptive field. (**i**) Histogram showing the distribution of multipeak scores in PoSub-HD and ADN-HD cells (Mann-Whitney U test, n = (1602, 97), Z = 4.25, *P* < 10^−4^). Shaded area, multi-peaked cells. Dotted lines, medians of the depicted distributions. (**j**) Average Fourier spectra of PoSub-FS cells (n = 427), PoSub-HD cells (n = 789) and ADN-HD cells (n = 69) after exclusion of multipeaked HD cells (2-way ANOVA, Fourier component by cell type interaction: F_(9,11547)_ = 4.22, *P* < 10^−4^). Inset: cumulative distribution of the data in the main panel. (**k**) Statistical distance between individual PoSub-HD cell or ADN-HD cell Fourier spectra and the average Fourier spectrum of the PoSub-FS cell population after exclusion of multi-peaked HD cells (n = (789, 69), Z = 10.4, *P* < 10^−24^). Dotted lines, medians of the depicted distributions. *D*_*KL*_, Kullback-Leibler divergence. (**l**) Box plots showing statistical distance between individual Fourier spectra and corresponding population averages after exclusion of multi-peaked HD cells (ANOVA, cell type, F_(2,1282)_ = 9.12, P < 10^−60^). Boxes represent the interquartile range (IQR) and whiskers represent minimum and maximum values that are not outliers. Outliers were defined as values more than 1.5 x IQR away from the top or bottom of the box and are not shown on the panel. Individual comparisons (Bonferroni correction) displayed on the panel. (**m**) Phases of individual Fourier components are correlated in PoSub-HD cells (circular correlation, Phase 1 vs Phase 2: *ρ*_cc_ = 0.51, *P* < 10^−20^; Phase 1 vs Phase 3: *ρ*_cc_ = 0.43, *P* < 10^−20^) and ADN-HD cells (circular correlation, Phase 1 vs Phase 2: *ρ*_cc_ = 0.26, *P* < 0.01; Phase 1 vs Phase 3: *ρ*_cc_ = 0.44, *P* < 10^−5^) but not in PoSub-FS cells (circular correlation, Phase 1 vs Phase 2: *ρ*_cc_ = 0.006, *P* = 0.91; Phase 1 vs Phase 3: *ρ*_cc_ = 0.06, *P* = 0.26). *ρ*_cc_, circular correlation coefficient. (**n**) Average Fourier spectra of PoSub-FS cells in experimental conditions shown in Fig. 1 (2-way ANOVA, Fourier component by cell type interaction. Left: n = 427, F_(9,3834)_ = 2.03, *P* = 0.10; Middle: n = 264, F_(9,2367)_ = 3.68, *P* < 0.05; Right: n = 99, F_(9,882)_ = 1.40, *P* = 0.24. Insets: histograms comparing symmetry scores (SymScores) of PoSub-FS cells in corresponding conditions (Wilcoxon signed rank test. Left: 1^st^ half vs 2^nd^ half, n = 427, Z = 1.13, *P* = 0.25; Middle: square vs triangle, n = 264, Z = 0.18, *P* = 0.86; Right: cue position 1 vs cue position 2, n = 99, Z = 0.02, *P* = 0.98).

**Extended Data Fig. 4.**
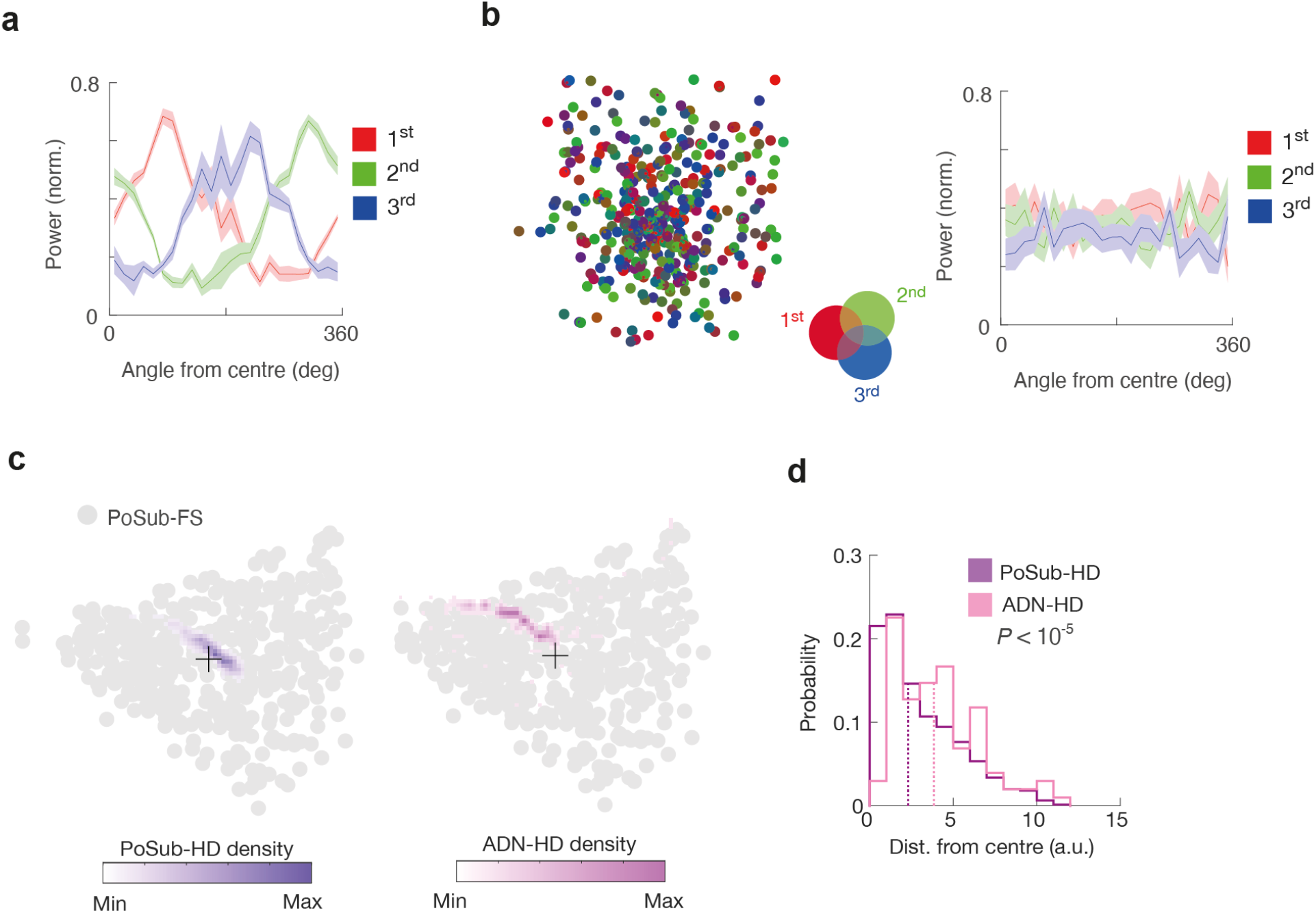
PoSub-FS cell tuning shows a continuous distribution of the first three Fourier components. (**a**) Relative power of the first three Fourier components as a function of the angular coordinate from the centre of the Isomap projection shown in **Fig. 2i**. Shaded lines represent mean +/- SEM. (**b**) Left, equivalent Isomap projection of time-reversed control PoSub-FS cell tuning curve auto-correlograms. Right, relative power of the first three Fourier components as a function of the angular coordinate from the centre of the control Isomap projection shown on the left. (**c**) Joint two-dimensional projection of PoSub-FS cell tuning curve auto-correlograms (gray points) and (left) PoSub-HD cell or (right) ADN-HD cell auto-correlograms. Values for PoSub-FS cell are shown as individual points (gray) underneath the density distribution of HD cell values. Densities below 0.1 of the maximum value are not shown. Black crosses indicate the centre of the PoSub-FS cell distribution. (**d**) Histogram showing Euclidean distance between individual PoSub-HD or ADN-HD cells and the centre of PoSub-FS distribution within the Isomap projections shown in (**c**) (n = (1602, 97), Mann-Whitney U, Z = 4.26 *P* < 10^−9^). Dotted lines show the medians of the depicted distributions.

**Extended Data Fig. 5.**
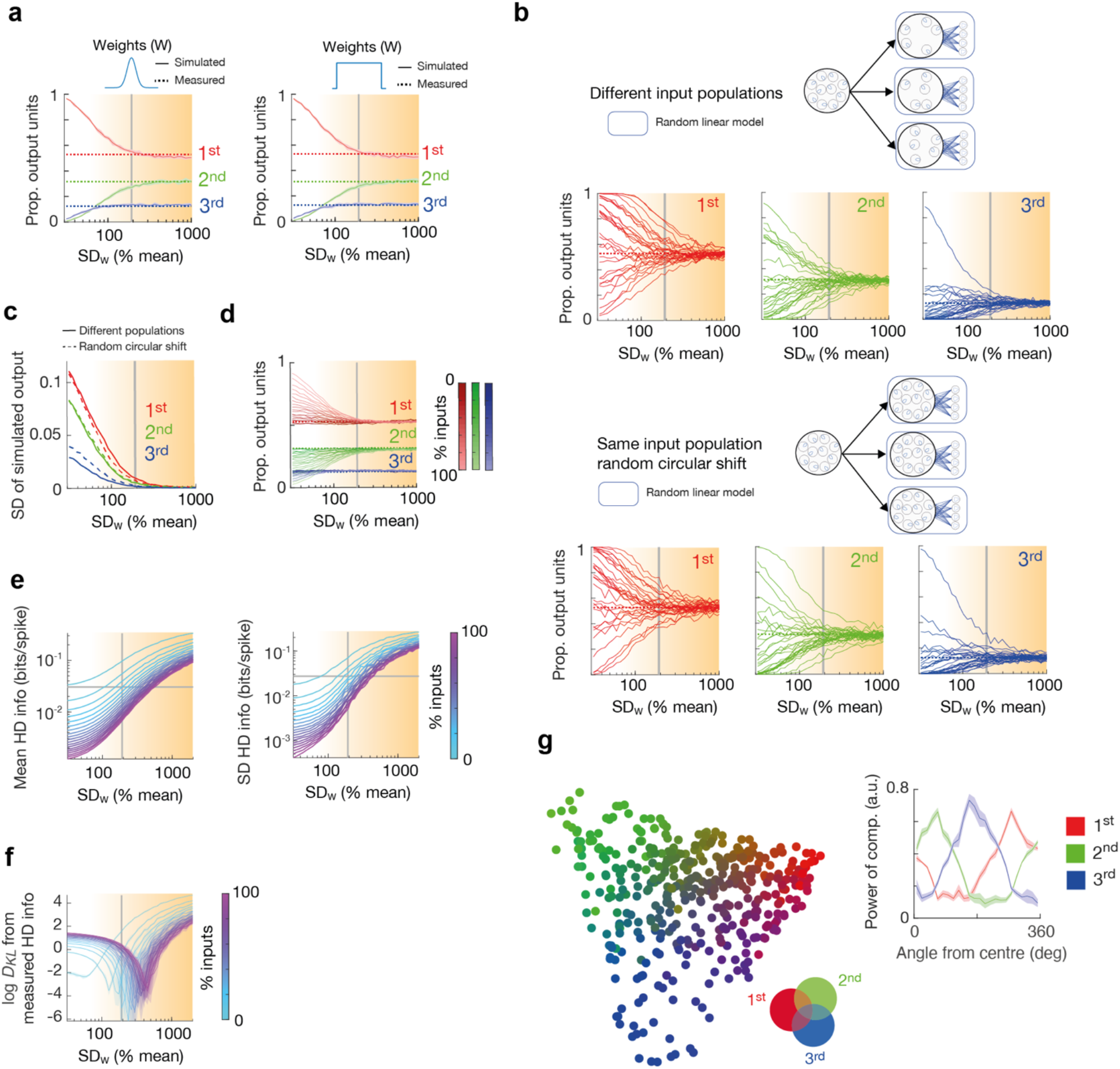
PoSub-FS like tuning curves emerge from random connectivity in a linear regime. (**a**) Proportion of simulated cells with maximum power in the first three Fourier components as a function of spread of synaptic weight distribution. Left: normal distribution, right: uniform distribution. Dotted lines display actual proportion of recorded PoSub-FS cells. Shaded area of each curve, SD based on 40 simulations. (**b**) Proportions of output units with the highest *n*th Fourier component across multiple simulations. Top, sampling from different sub-populations of the pool of input tuning curves. Bottom, sampling from the whole pool of input tuning curves with a random circular shift. (**c**) Quantification of variance across individual simulations shown in (**b**). (**d**) Proportion of output unit with the highest *n*th Fourier component as a function of connection sparsity (percentage of inputs shared between output units). (**e**) Mean and standard deviation (SD) of HD information of simulated output tuning curves as a function of synaptic weight distribution and input sparsity. (**f**) Difference in HD information distribution between FS-PoSub cell tuning curves and simulated output tuning curves as a function of synaptic weight distribution and sparsity of connectivity. *D*_*KL*_, Kullback-Leibler divergence. (**g**) Left, Isomap projection of simulated output tuning curve auto-correlograms. Right, relative power of the first three Fourier components as a function of the angular coordinate from the centre of the Isomap projection shown on the left.

**Extended Data Fig. 6.**
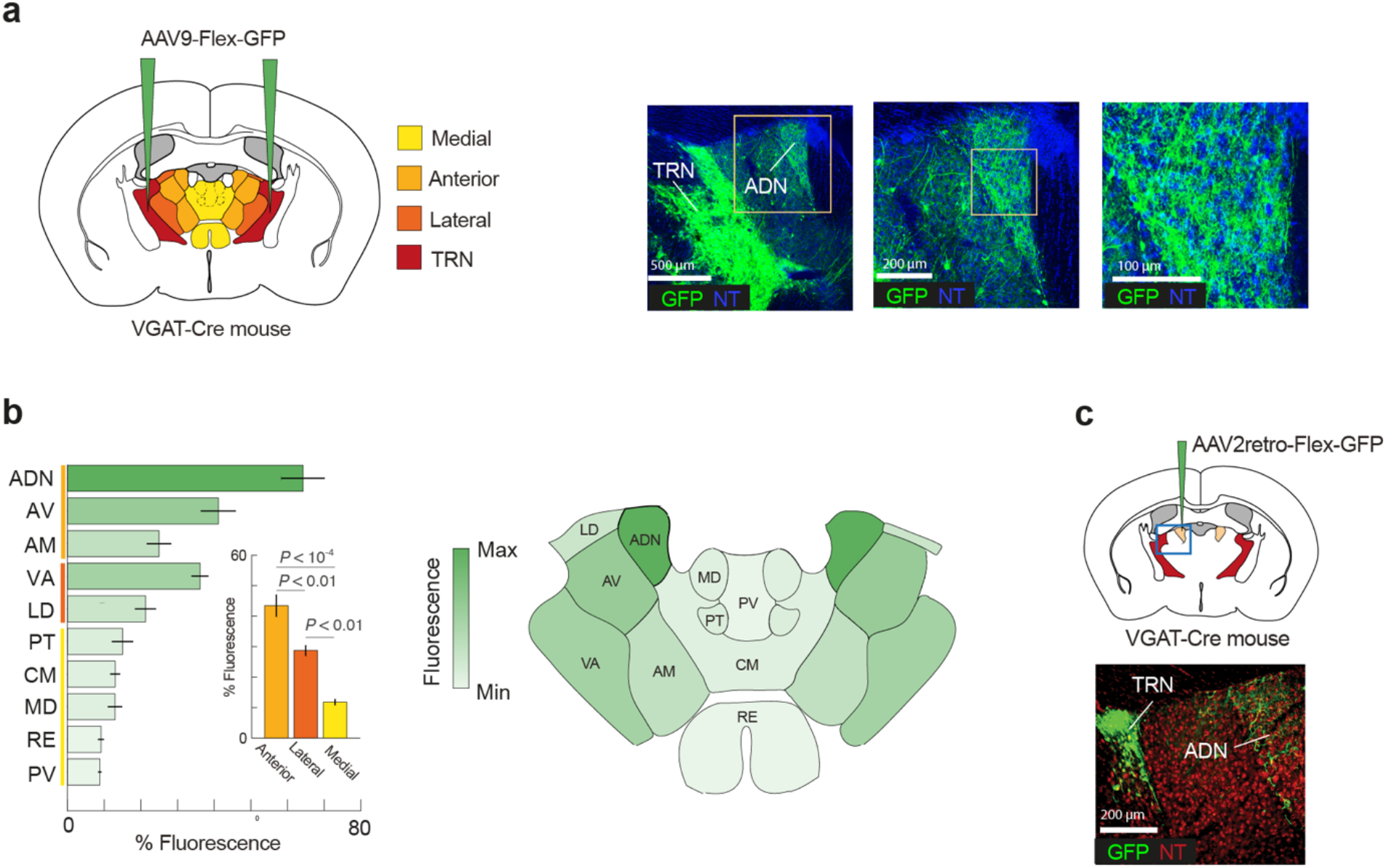
ADN is densely innervated by inhibitory afferents from the TRN. (**a**-**b**), Anterograde tracing of projections from TRN to the rostral thalamus. (**a**) Left: brain diagram showing virus injection sites and sub-divisions of the rostral thalamic nuclei. Right: representative images of fluorescent signal in TRN and neighboring thalamic nuclei. (**b**) Left: TRN innervation density in rostral thalamic nuclei (n = 4 mice). Left, inset: density of TRN projections across sub-divisions of the rostral thalamus (repeated measures ANOVA, F_(2,9)_ = 44.9, *P* < 10^−4^). Individual comparisons (Bonferroni correction) displayed on the panel. Data shown as mean +/- SEM. Right: individual rostral thalamic nuclei colour-coded according to the TRN projection strength. ADN, anterodorsal nucleus; AV, anteroventral nucleus; AM, anteromedial nucleus; VA, ventral anterior nucleus; LD, laterodorsal nucleus; PT, paratenial nucleus; CM, centromedial nucleus; MD, mediodorsal nucleus; RE, nucleus reuniens; PV, paraventricular nucleus. (**c**) Retrograde tracing of projections from TRN to ADN (n = 2 replicates). Top: coronal diagram showing the virus injection site. Bottom: representative image of fluorescent signal in ADN and anterodorsal TRN.

**Extended Data Fig. 7.**
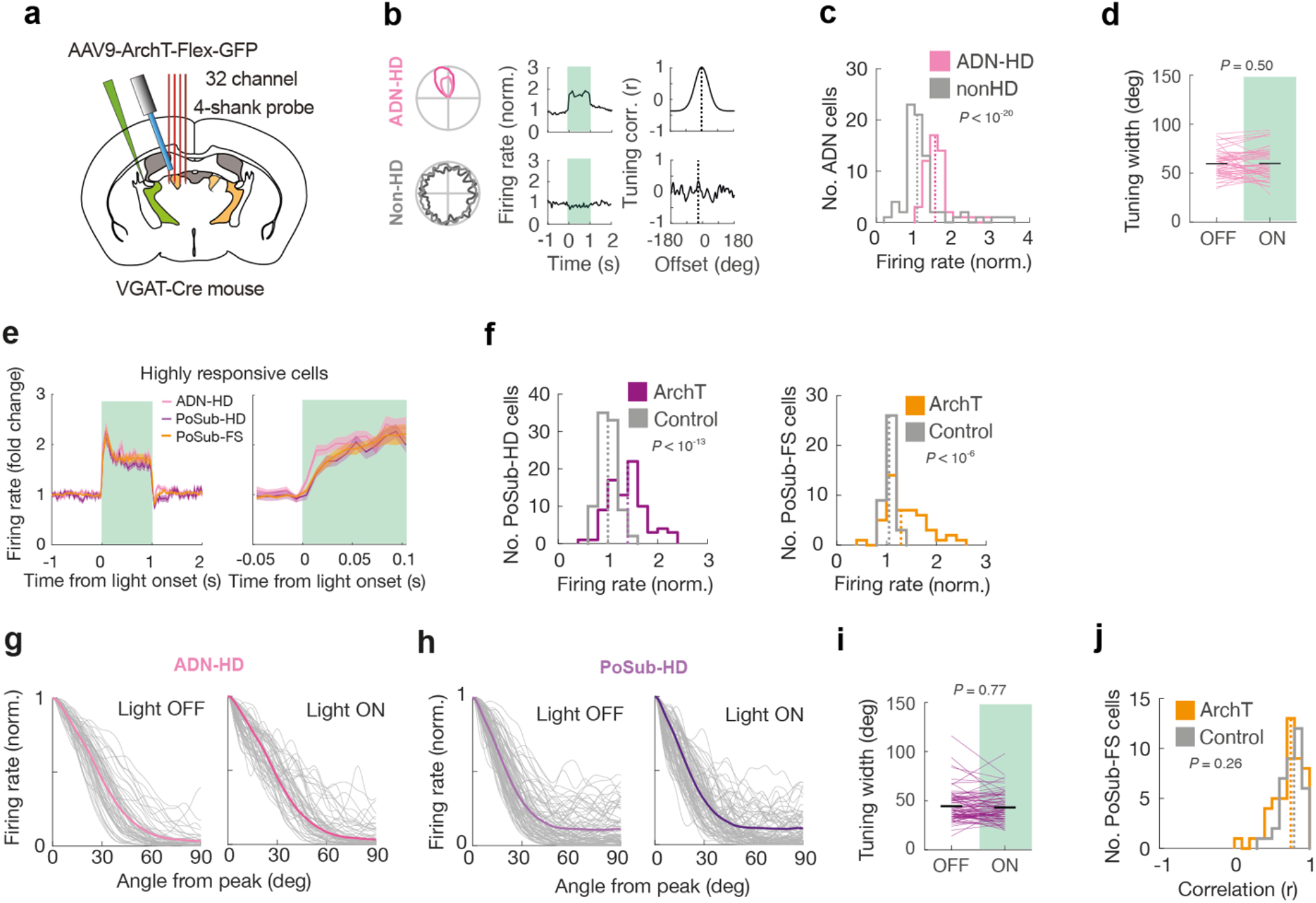
Disinhibition of ADN-HD cells and comparison with other cell populations. (**a**-**b**) Selective optogenetic modulation of thalamic HD gain. (**a**) Brain diagram of the unilateral viral injection into anterior TRN and positioning of the probe and optic fiber above ADN. (**b**) Representative examples of ADN-HD cell and thalamic non-HD cell responses to optogenetic inactivation of TRN projections to ADN. Left: HD tuning curves during light OFF epoch (light shades) and light ON epoch (dark shades). Middle: Average effect of the optogenetic manipulation on representative cells’ firing rates. Green shading represents the duration of the light pulse. Right: cross-correlation between HD tuning curves during the light ON and light OFF epochs. (**c**) Histograms of firing rate modulation in ADN-HD and Non-HD cells (n = (52, 75), Mann-Whitney U test between cell types, Z = 6.57 *P* < 10^−20^). Dotted lines show the medians of the depicted distributions. (**d**) Width of individual ADN-HD cell tuning curves in light ON and light OFF epochs (n = 52; Wilcoxon signed rank test, Z = 0.67, *P* = 0.50). Horizontal lines represent medians of each condition. (**e**) Temporal profile of highly-responsive ADN-HD, PoSub-HD and PoSub-FS responses to optogenetic ADN disinhibition. Only cells with average response above the median for a given population were included (n = 26, 41, 23). (**f**) Histograms of firing rate modulation in PoSub-HD cells (top; ArchT, n = 83; Control: n = 89; Mann-Whitney U test vs control group, Z = 1.69, *P* < 10^−13^) and PoSub-FS cells (bottom; ArchT, n = 47; Control: n = 38; Mann-Whitney U test vs control group, Z = 4.99, *P* < 10^−6^). Dotted lines, medians of the depicted distributions. (**g**-**h**) Tuning curve slopes of ADN-HD cells (n = 52) and PoSub-HD cells (n = 83) during light ON and light OFF epochs. Thin gray lines denote individual tuning curves, thick colored lines denote population averages. (**g**) Width of individual PoSub-HD cell tuning curves in light ON and light OFF epochs (n = 52; Wilcoxon signed rank test, Z = 0.30, *P* = 0.50). Horizontal lines, medians of each condition. (**h**) Correlation between PoSub-FS cell tuning curves during light ON and light OFF epochs (ArchT group, n = 47; Control group: n = 38; Mann-Whitney U test, Z = 1.14, *P* = 0.26). Dotted lines show the medians of the depicted distributions.

**Extended Data Fig. 8.**
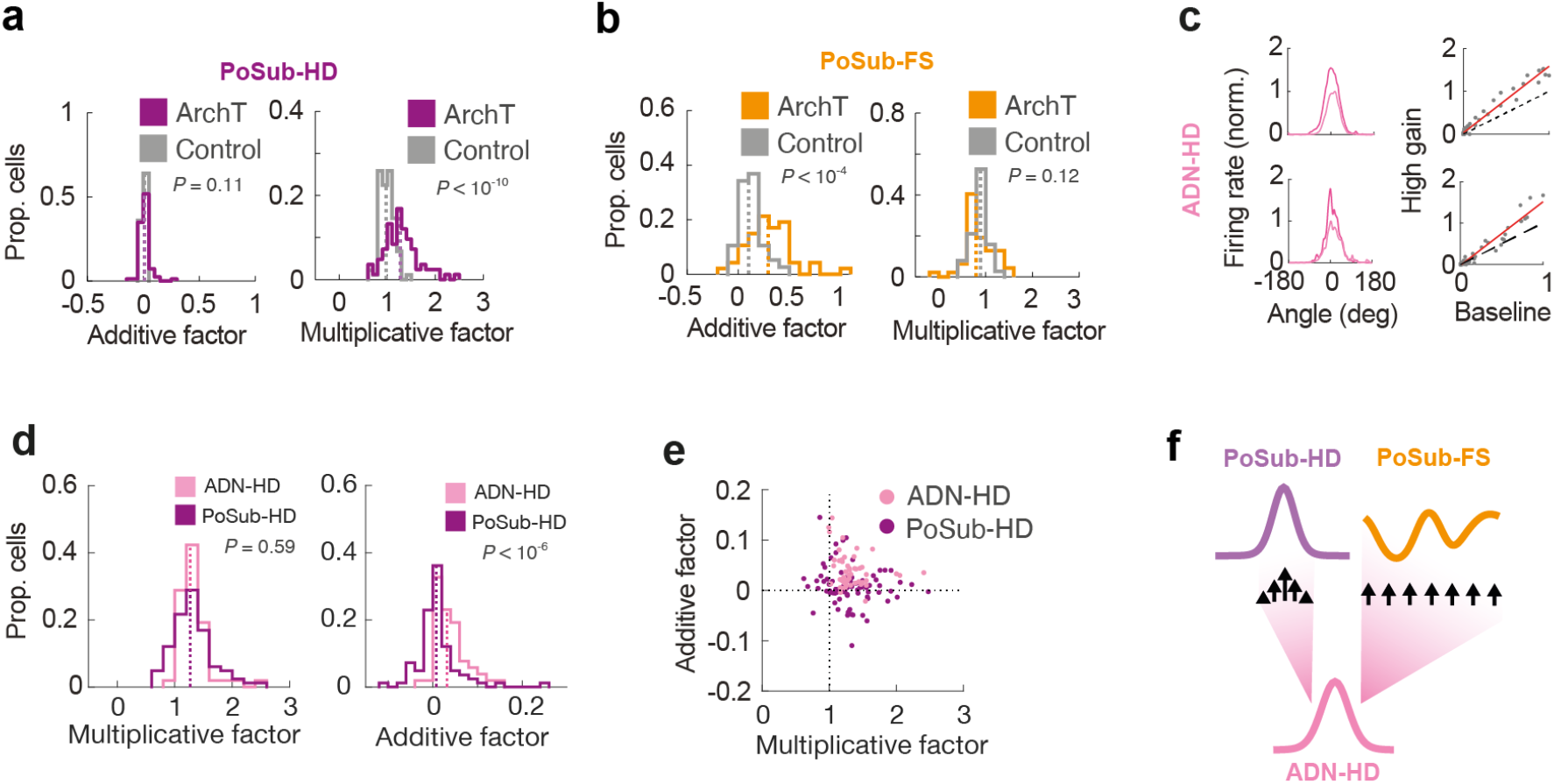
PoSub-HD cells exhibit only multiplicative gain, PoSub-FS cells only additive gain, and ADN-HD cells a mixture of both. (**a**) Histograms of of additive and multiplicative factors for PoSub-HD cells (Additive factor: Mann-Whitney U test vs Control group, Z = 1.56, *P* = 0.11; Multiplicative factor: Mann-Whitney U test vs Control group, Z = 6.80, *P* < 10^−10^). Dotted lines show the medians of the depicted distributions. (**b**) Histograms of additive and multiplicative factors for PoSub-FS cells (Additive factor: Mann-Whitney U test vs Control group, Z = 4.26, *P* < 10^−6^; Multiplicative factor: Mann-Whitney U test vs Control group, Z = 0.53, *P* = 0.12). Dotted lines represent the medians of the depicted distributions. (**c**) Representative examples of ADN-HD cells in baseline condition (light shades) and high gain condition (dark shades) plotted in cartesian coordinates and their respective tuning correlation plots. Red lines represent the linear fit. (**d**) Histograms comparing ADN-HD and PoSub-HD cell populations in terms of multiplicative factors (Mann-Whitney U test, Z = 0.55, *P* = 0.58) and additive factors (Mann-Whitney U test, Z = 5.07, *P* < 10^−6^). Dotted lines show the medians of the depicted populations. (**e**) Scatter plot showing relative contributions of additive and multiplicative factors in PoSub-HD and ADN-HD cells.

**Extended Data Fig. 9.**
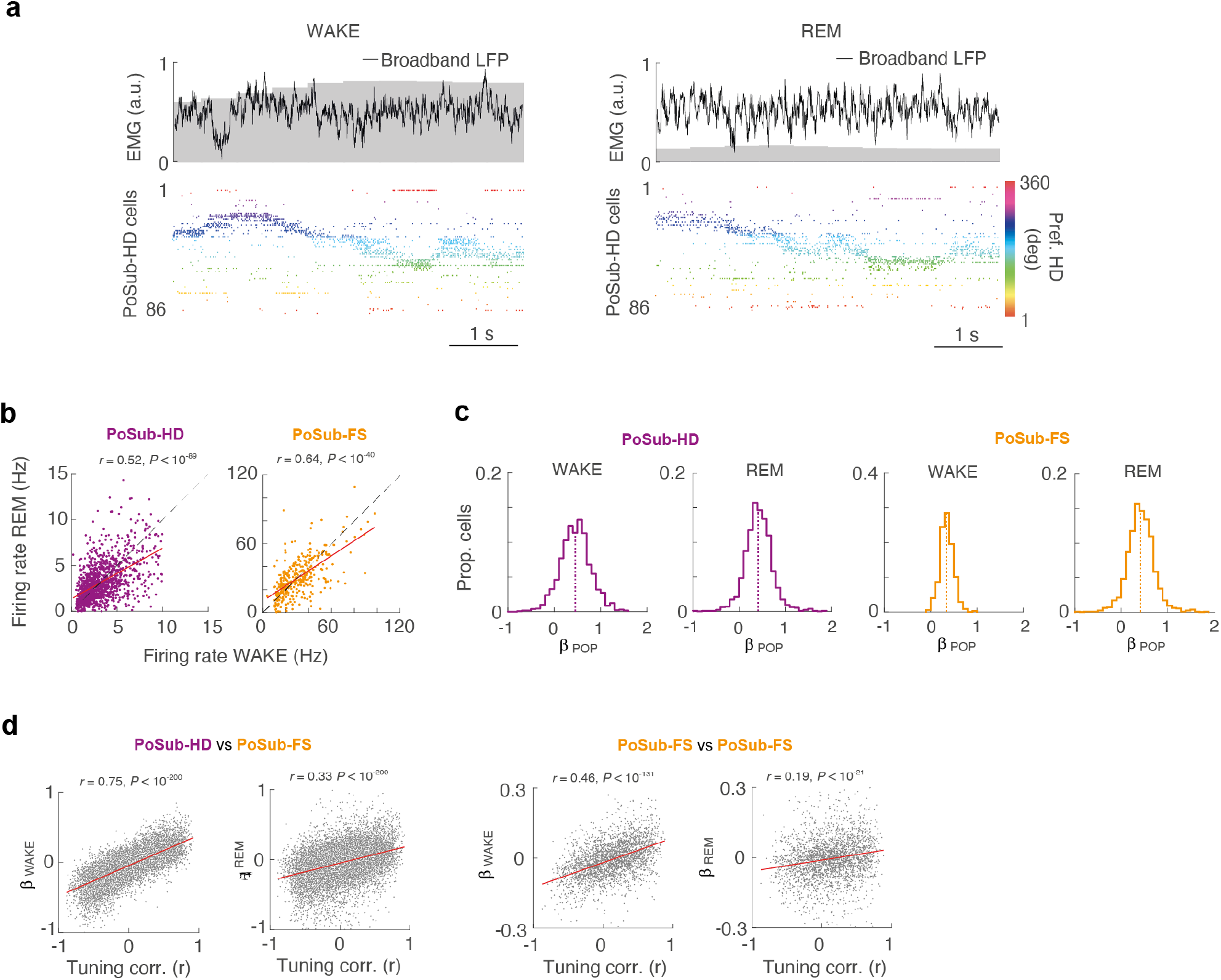
Single-unit spiking characteristics, population coupling and coupling-tuning relationship during WAKE and REM. (**a**) Representative LFP traces and PoSub-HD cell raster plots during WAKE and REM epochs. Grey areas indicate level of LFP-derived electromyogram (EMG, see Methods). HD cells are sorted and color-coded according to their preferred direction. (**b**) Scatter plots of firing rates during WAKE and REM epochs for PoSub-HD and PoSub-FS cells. (**c**) Histograms of GLM coupling of individual cells to the population firing rate. β, cross-coupling coefficient. (**d**) Scatter plots showing the relationship between HD tuning correlation and GLM pairwise coupling for PoSub-HD:PoSub-FS and PoSub-FS:PoSub-FS cell pairs during WAKE and REM. β, cross-coupling coefficient.

**Extended Data Fig. 10.**
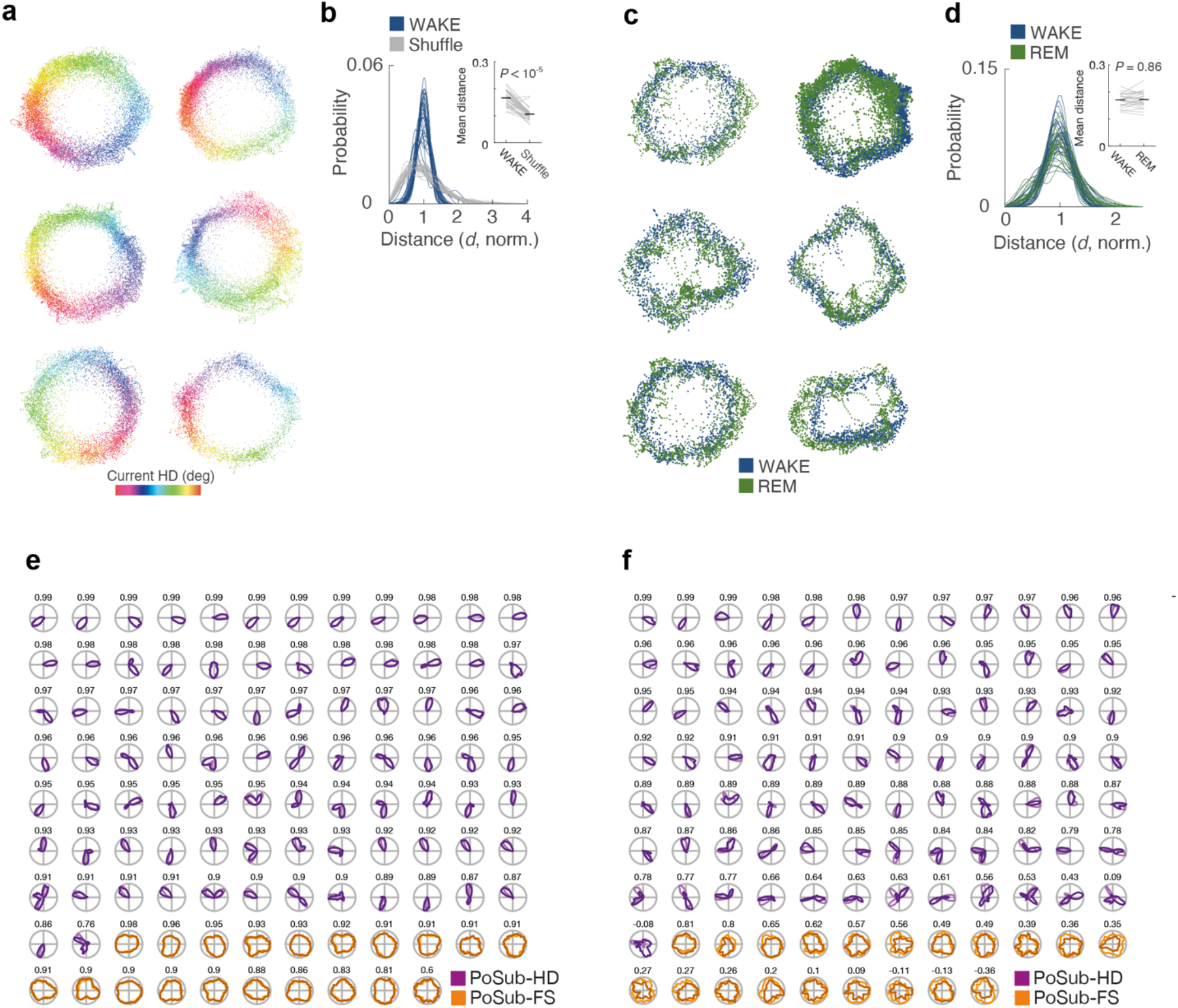
Isomap projections during WAKE and REM and additional tuning curve examples. (**a**) Additional examples of Isomap projections of HD population vectors during WAKE from six recordings. (**b**) Distributions of distance values to the centre of the manifold for all WAKE Isomap projections of real and shuffled HD population vectors, normalized to the mean distance of the real projections. Each curve represents one recording session. Inset: mean distance to the centre of the manifold for real Isomap projections (n = 32 mice; Wilcoxon signed rank test vs shuffle, Z = 4.79, *P* < 10^−5^). (**c**) Real HD tuning curves (light shades) and WAKE Isomap HD tuning curves (dark shades) of all PoSub-HD and PoSub-FS cells from a single recording (same as in **Fig. 6B**). Pearson correlation coefficients depicted above each pair of tuning curves. (**d**) Additional examples of Isomap projections of HD population vectors during WAKE and REM from six recordings (same as in **A**). Pearson correlation coefficients depicted above each pair of tuning curves. (**e**) Distributions of distance values to the centre of the manifold for all REM and subsampled WAKE Isomap projections, normalized to the mean distance of the WAKE projections. Each curve represents one recording session. Inset: mean distance to the centre of the manifold for REM and subsampled WAKE Isomap projections (n = 26 mice; Wilcoxon signed rank test, Z = 0.17, *P* = 0.86). (**f**) WAKE (light shades) and REM (dark shades) Isomap tuning curves of all PoSub-HD and PoSub-FS cells from a single recording (same as in **Fig. 6e**).

